# Sex-specific transcription and DNA methylation landscapes of the Asian citrus psyllid, a vector of huanglongbing pathogens

**DOI:** 10.1101/2022.01.28.478167

**Authors:** Xiudao Yu, Hollie Marshall, Yan Liu, Yu Xiong, Xiangdong Zeng, Haizhong Yu, Wei Chen, Guchun Zhou, Bo Zhu, Laura Ross, Zhanjun Lu

## Abstract

The relationship of DNA methylation and sex-biased gene expression is of high interest, it allows research into mechanisms of sexual dimorphism and the development of potential novel strategies for insect pest control. The Asian citrus psyllid, Diaphorina citri Kuwayama, is a major vector for the causative agents of Huanglongbing (HLB), which presents an unparalleled challenge to citrus production worldwide. Here, we identify the X chromosome of D. citri and investigate differences in the transcription and DNA methylation landscapes between adult virgin males and females. We find a large number of male-biased genes on the autosomes and a depletion of such on the X chromosome. We have also characterised the methylome of D. citri, finding low genome-wide levels, which is unusual for an hemipteran species, as well as evidence for both promoter and TE methylation. Overall, DNA methylation profiles are similar between the sexes but with a small number of differentially methylated genes found to be involved in sex differentiation. There also appears to be no direct relationship between differential DNA methylation and differential gene expression. Our findings lay the groundwork for the development of novel epigenetic-based pest control methods, and given the similarity of the D. citri methylome to other insect species, these methods could be applicable across agricultural insect pests.

## Introduction

Sexual dimorphisms in behavior, morphology and physiology are widespread across sexually reproducing organisms, oftentimes producing dramatic phenotypic differences between sexes. Sexual dimorphisms between males and females have been associated with a number of genomic processes across taxa such as: genetic differences e.g. sex chromosomes (Mank, 2009), alternative splicing (Wexler *et al*., 2019), genomic imprinting (Zou *et al*., 2020) and epigenetic mechanisms (Bain *et al*., 2021). These mechanisms result in sex-biased gene expression, which is generally thought to underlie most sex-specific differentiation (Ledón-Rettig *et al*., 2017).

DNA methylation (one of the most well-studied and conserved epigenetic modifications) has recently been shown to be important in sexual dimorphism in some hemipteran insects (Bain *et al*., 2021). DNA methylation in insects refers synonymously with cytosine methylation in a CG dinucleotide (CpG) context, with very little cytosine methylation reported in a non-CpG context in insects (Bonasio *et al*., 2012). In comparison with mammals and plants, insect genomes have sparse cytosine methylation mainly restricted to exons of transcribed genes (although see Lewis *et al*. (2020)), with typically less than 3% of cytosines methylated (Glastad *et al*., 2019). Some holometabolous insects are even reported to have no detectable levels of DNA methylation—e.g., the coleopteran *Tribolium castaneum* and dipteran *Drosophila* (Bewick *et al*., 2017; Zemach and Zilberman, 2010). DNA methylation in arthropods is preferentially targeted to genes that perform core and conserved “housekeeping” functions (Bewick *et al*., 2017; Glastad *et al*., 2019). Within these genes, DNA methylation is largely confined within coding regions of holometabolous insects (Bewick *et al*., 2017; Lewis *et al*., 2020); whereas hemimetabolous insects have a relatively higher and more global methylation, in which DNA methylation extends to the introns, transposable elements (TEs) and gene promoters (Bain *et al*., 2021; Lewis *et al*., 2020), as is also seen in Blattodea (Bewick *et al*., 2017), and Orthoptera (Falckenhayn *et al*., 2013; Lo *et al*., 2018).

DNA methylation aids in genome stability and proper regulation of gene expression for many species. In vertebrates and plants, CpG methylation functions for genome defence against invading TEs, once recognized by the host, TEs would be methylated, thereby preventing their transcription and transposition, and loss of DNA methylation leads to reactivation of TEs (Glastad *et al*., 2019; Walsh *et al*., 1998; Zemach and Zilberman, 2010). Associations between DNA methylation and transcriptional regulation usually depend upon genomic context. DNA methylation of regulatory elements in vertebrates e.g. gene promoters, can suppress gene expression levels by affecting transcription factor binding and recruitment of the transcription initiation complex, typically causing stable silencing. Whereas gene-body DNA methylation, as is found in insects is generally reported to be associated with elevated, stable gene expression (Cardoso-Júnior *et al*., 2021; Glastad *et al*., 2016; Libbrecht *et al*., 2016). This may attribute to the fact that DNA methylomes profiled to date in insects have retained CpG methylation within constitutivly expressed gene bodies, which is proposed to affect gene expression through regulation of transcription elongation and alternative splicing (Bewick *et al*., 2017; Bonasio *et al*., 2012; Lewis *et al*., 2020; Lorincz *et al*., 2004). Accordingly, former studies documented that intragenic DNA methylation could improve the transcriptional fidelity in mice (Neri *et al*., 2017), as well as enhance the expression level of genes in *Arabidopsis* (Shahzad *et al*., 2021).

Gene expression largely contributes to the sexual and morphological differences between insect morphs. Several researchers have sought to characterize regulatory mechanisms e.g. DNA methylation, that may govern sex-specific gene expression in insects. Sex-specific patterns of DNA methylation that may be implicated in sexual dimorphisms has been studied in the hemipteran peach aphid *Myzus persicae* and the citrus mealybug *Planococcus citri* (Bain *et al*., 2021; Mathers *et al*., 2019). By way of example in *M. persicae*, about 19% of genes are differentially expressed between sexes, exhibiting a positive correlation between changes in gene expression and DNA methylation, where the higher sex-specific gene expression is usually accompanied with higher sex-specific methylation, particularly for the genes located on sex chromosomes (Mathers *et al*., 2019). Strikingly, both sexual dimorphisms and paternal genome elimination in *P. citri*, an insect that has no sex chromosomes and displays a unique pattern of DNA methylation with the presence of promoter methylation, are predicted to be under epigenetic control; however the *cis*-acting DNA methylation profiles fail to explain its sex-biased gene expression patterns (Bain *et al*., 2021). Therefore, despite some evidence suggesting a relationship between DNA methylation and gene expression, whether or not changes in CpG methylation directly regulate sex-biased gene expression need to be explored in other insect species.

The Asian citrus psyllid, *Diaphorina citri* Kuwayama, is the most important citrus pest because it efficiently transmits *Candidatus Liberibacter asiaticus* (CLas) pathogen, which causes citrus huanglongbing disease in most citrus-producing regions of the world (Grafton-Cardwell *et al*., 2013; Yu and Killiny, 2020). Development of novel genetic control techniques (e.g. RNAi strategy and sterile insect technique) against *D. citri* requires a specific understanding of the molecular mechanisms governing key physiological traits, such as sexual dimorphisms. Males and females of *D. citri*, in the absence of confounding genetic variation, are phenotypically indistinguishable before emerging as adults (Yu and Killiny, 2018). Here, we identify genome-wide differences in DNA methylation between male and female *D. citri*, including the relationship of DNA methylation with gene expression and the DNA methylation differences in the sex chromosomes. We performed investigations as follows: (i) an identification of the X chromosome; (ii) a study of the sex-specific DNA methylation landscape; (iii) an analysis of sex-biased gene expression and DNA methylation; and (iv) a genome-wide comparison between DNA methylation and gene expression.

## Methods

### Insect rearing

A *D. citri* colony was continuously reared at the National Navel Orange Engineering Research Center, Gannan Normal University, Jiangxi, China. The culture was established in 2015 using field populations from Nankang District, Jiangxi and maintained on *Murraya exotica* seedlings in a greenhouse set at 27 °C ± 1°C and relative humidity of 70% ± 5% with a 14:10h light:dark photoperiod. Newly emerged adults were collected for sex separation under a stereomicroscope, and then kept in the separated cages with new *M. exotica* seedlings.

### RNA and DNA extraction and sequencing

Groups of twenty 3-day virgin females or males for DNA/RNA extraction were collected between 3 p.m. and 4 p.m. every day to avoid differences due to the circadian rhythm (Pegoraro *et al*., 2016). RNA extraction was performed using an RNeasy Kit (QIAGEN, Valencia, CA, USA), and genomic DNA was extracted using the Qiagen DNeasy Blood Tissue Kit (QIAGEN, Valencia, CA, USA), according to the manufacturer’s instructions. RNA/DNA degradation and contamination was validated on 1% agarose gels. The concentration was measured using Qubit^®^ DNA Assay Kit in Qubit^®^ 2.0 Flurometer (Life Technologies, Carlsbad, CA, USA). An amount of 100 ng genomic DNA (for whole genome sequencing - WGS) or 100 ng genomic DNA spiked with 0.5 ng lambda DNA (for whole genome bisulfite sequencing - WGBS) were fragmented by sonication to 200-300 bp with Covaris S220 (Covaris Inc., Woburn, MA, USA). For WGBS, the fragmentized DNA samples were treated with bisulfite using EZ DNA Methylation-GoldTM Kit (Zymo Research, Irvine, CA, USA). For RNA-seq, a total of 3.0*μ*g RNA per sample was used to prepare the sequencing libraries using the NEBNext UltraTM RNA Library Prep Kit for Illumina (NEB, Ipswich, MA, USA). All above libraries were constructed and sequenced by Novogene Corporation (Beijing, China). 150bp paired-end sequencing of each sample was performed on an Illumina NovaSeq 6000 platform (Illumina Inc., San Diego, CA, USA). Library quality was assessed on the Agilent Bioanalyzer 2100 system (Agilent Technologies, Santa Clara, CA, US).

### X chromosome identification

Whole genome sequencing of a pool of males and a pool of females was used to identify the X chromosome. Data were quality checked using Fastqc v0.11.8 (Andrews, 2010) and aligned to the reference genome (Diaci v3.0, Hosmani *et al*. (2019)) using Bowtie2 v2.3.5.1 (Langmead and Salzberg, 2013) in *sensitive* mode. Coverage per chromosome was then calculated using samtools v.1.9 (Li *et al*., 2009). Coverage levels were normalised by chromosome length and mean coverage per sample and the log2 ratio of male to female coverage was plotted using R v4.0.3 (R Core Team, 2020). We also repeated the above analysis using 10,000bp windows across each chromosome to check for any incorrectly assembled X-linked regions.

To provide further evidence for the identification of the X chromosome, we carried out a synteny analysis between *D. citri* and the psyllid *Pachypsylla venusta* (genome: Pven_dovetail (Li *et al*., 2020)). The protein sequences of single copy genes from *D. citri* were blasted against those from *P. venusta* using blastp v2.2.31 (Camacho *et al*., 2009) with an e-value of 1e^-10^. MCScanX (Wang *et al*., 2012) was then used to identify coliniarity blocks across genomes using the interspecies setting and requiring a minimum of 10 genes per block. Results were visualised using Synvisio (Bandi and Gutwin, 2020).

### Differential gene expression

RNA-seq was carried out on pools of males and females with three replicates per sex. Data were quality checked using Fastqc v0.11.8 (Andrews, 2010) and quality trimmed using CutAdapt v1.18 (Martin, 2011). Reads were aligned to the reference genome (Diaci v3.0 (Hosmani *et al*., 2019)) and transcript abundances were calculated using RSEM v1.3.3 (Li and Dewey, 2011) implementing STAR v2.7.3a (Dobin *et al*., 2013). DESeq2 v1.28.1 (Love *et al*., 2014) was used to determine differentially expressed genes between males and females. A gene was considered differentially expressed if the corrected p-value was <0.05 (adjusted for multiple testing using the Benjamini-Hochberg procedure (Benjamini and Hochberg, 1995)) and the log2 fold-change was >1.5. Chromosome enrichment of sex-biased genes was determined using the hypergeometric text implemented in R v4.0.3 (R Core Team, 2020).

### Genome-wide DNA methylation and differential DNA methylation

Whole genome bisulfite sequencing data were quality checked using Fastqc v0.11.8 (Andrews, 2010) and reads were aligned to the reference genome (Diaci v3.0 (Hosmani *et al*., 2019)) using Bismark v0.20.0 (Krueger and Andrews, 2011). Reads were also aligned to the *E. coli* phage lambda reference (NCBI Accession: PRJNA485481) in order to determine the bisulfite conversion efficiency. Weighted methylation levels of genomic features were calculated as in Schultz *et al*. (2012).

Differentially methylated CpG sites were determined using the R package MethylKit v1.16.0 (Akalin *et al*., 2012). Coverage outliers (above the 99.9 percentile) and bases with <10 coverage were removed. A binomial test was then carried out per CpG position per sample using the lambda conversion rate as the probability of success and correcting P-values using the Benjamini-Hochberg procedure (Benjamini and Hochberg, 1995). Only sites which were classified as methylated in at least one sample were used for final differential methylation analysis. A logistic regression model implemented by MethylKit (Akalin *et al*., 2012) was then used to determine differentially methylated CpGs between sexes. A site was classified as differentially methylated if the Benjamini-Hochberg (Benjamini and Hochberg, 1995) corrected P-value <0.01 and the overall methylation difference was >10%. Exon regions were classed as differentially methylated if they contained at least two differentially methylated CpG and had an overall weighted methylation difference of >15%. Two CpGs were chosen as Xu *et al*. (2021) found methylation of two CpGs within a region were enough to induce gene expression changes via histone recruitment in the silk moth.

### Relationship between gene expression and DNA methylation

The relationship between DNA methylation and gene expression was determined using linear models implemented by custom scripts in R (R Core Team, 2020). Interaction effects were determined using the *anova* function and post-hoc testing of fixed factors was done using the *glht* function from the multcomp R package (Hothorn *et al*., 2008) with correction for multiple testing by the single-step method. Correlation were calculated using Spearman’s rank correlation *rho*.

### Additional genome annotation and gene ontology enrichment

Transposable elements were *de novo* annotated in the *D. citri* genome using the EDTA pipeline (Ou *et al*., 2019). Putative promoter regions were defined as 500bp upstream of UTR regions. We excluded any promoters which overlap with other genomic features. Intergenic regions were determined as regions between gene end and gene start sites (excluding the newly annotated putative promoters and excluding any TE overlap).

Additional gene ontology annotations were generated from the protein sequences of all genes using eggNOG-mapper v.2.0.0 with standard parameters (Cantalapiedra *et al*., 2021). A total of 14,133 genes were annotated with GO terms. GO enrichment was carried out in R using GOstats v2.56.0 (Falcon and Gentleman, 2007) which implements a hypergeometric test with Benjamini–Hochberg correction for multiple testing (Benjamini and Hochberg, 1995). GO biological processes were classes as over represented if the correct P-value was <0.05. REVIGO (Supek *et al*., 2011) was then used to visualise GO terms. GO terms for genes with high DNA methylation and differentially methylated genes between the sexes were tested against all methylated genes as a background. Hypermethylated genes per sex were tested against a background of all differentially methylated genes. Differentially expressed genes were tested against a background of all genes present in the RNA-seq data with detectable expression, >10 FPKM in at least one sample. Over-expressed genes per sex were tested against a background of all differentially expressed genes.

## Results

### X chromosome identification

Whole genome sequencing of a pool of males and a pool of females was used to identify the X chromosome of *D. citri*. Around 80% of reads mapped to the *D. citri* reference genome (supplementary 1.0.0) which resulted in 50x coverage for the female sample and 51X coverage for the male sample. Most psyllid species possess an XO sex determination system (Riemann, 1966; Maryańska-nadachowska *et al*., 2014), where females carry two X chromosomes and males carry a single X and no Y. Using a coverage-based analysis we have found that chromosome 08 shows roughly half coverage in males compared to females (Fig.1a and 1b), indicating this is likely the X chromosome. We confirm this by showing high synteny between chromosome 08 of *D. citri* and the related psyllid *P. venusta* X chromosome (Fig.1c). Finally, as the reference genome is primarily based on male data we were able to search for a divergent Y chromosome. We checked the coverage ratio of 10,000bp windows across the genome, finding no clear peaks with a log2 male to female coverage ratio greater than 0.5 (supplementary Fig.S1) which would indicate higher coverage in males compared to females, this suggests there is no Y chromosome in *D. citri*.

**Figure 1:**
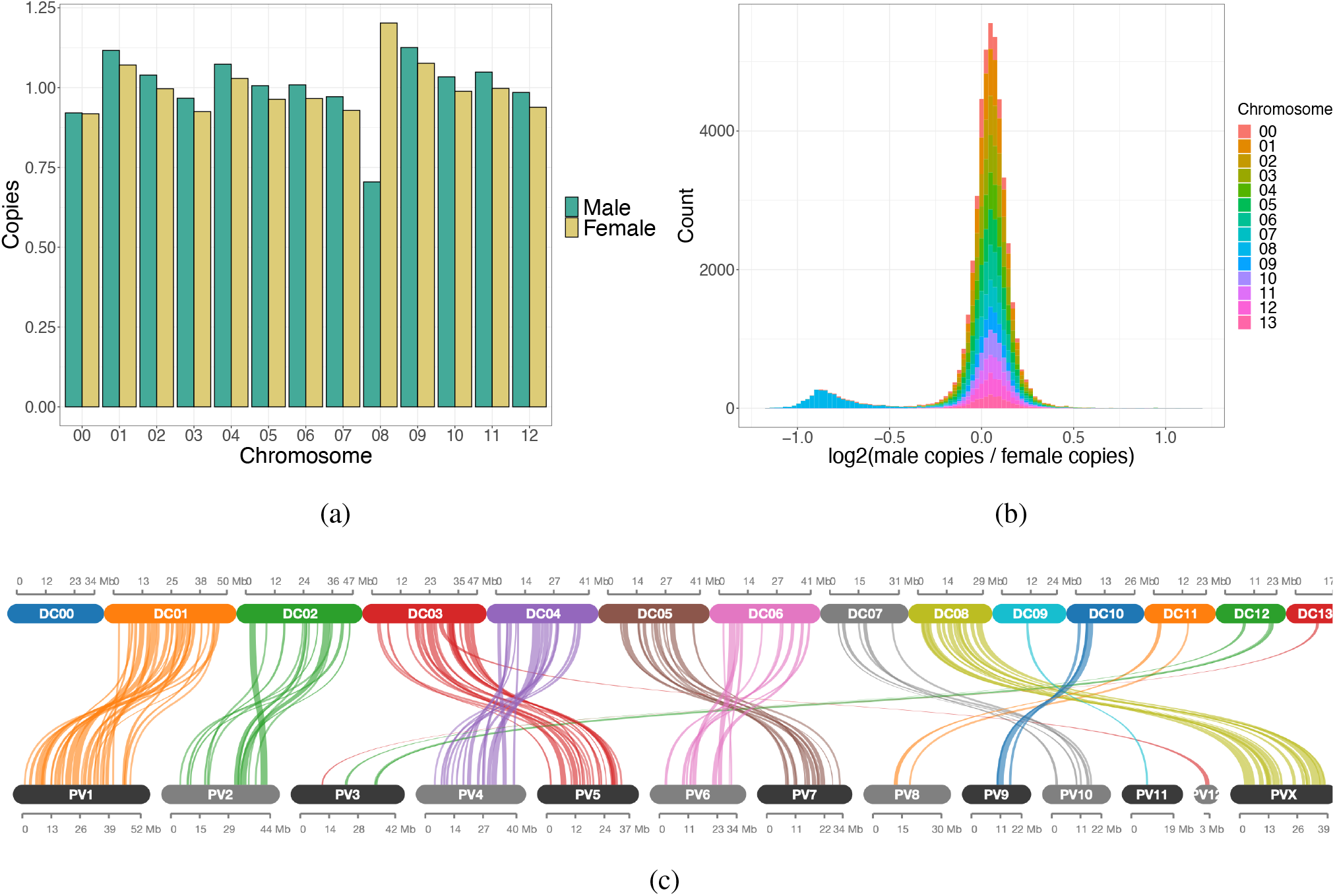
Identification of the X chromosome. (a) Bar plot of the coverage per chromosome for males and females normalised by the genome-wide average coverage. Chromosome 13 is missing from this graph as it represents unplaced scaffolds. (b) Histogram of the log2 male to female coverage ratio for 10,000bp windows across each chromosome. (c) Synteny plot showing collinearity blocks between *D. citri* (DC) and *P. venusta* (PV). Each line represents at least 10 orthologous genes.

### Sex-biased gene expression

Using RNA-seq from pools of males and females we have identified differentially expressed genes between the sexes. The majority of variation within the data (97%) is caused by sex (Fig. 2a) and whilst we find many genes have equal expression in both sexes, a large number of genes are only expressed in males (Fig. 2b). In total we have identified 1,259 genes out of 12,420 which are differentially expressed (adjusted P-value <0.05 and log2 fold-change >1.5, supplementary 1.0.1 and supplementary Fig.S2). Of these, significantly more are upregulated in males compared to females (Chi-squared goodness of fit: X-squared = 907.67, df = 1, P-value < 0.01). 1,164 are upregulated in males (9.4% of all genes tested) and 95 are upregulated in females (0.8% of all genes tested). A large number of the genes upregulated in males are also sex-limited (484 total, 41.6% of all male upregulated genes), meaning they have zero expression in females. Whilst only 12/95 genes (12.6%) upregulated in females are sex-limited (Fig.2c and Fig.2d).

**Figure 2:**
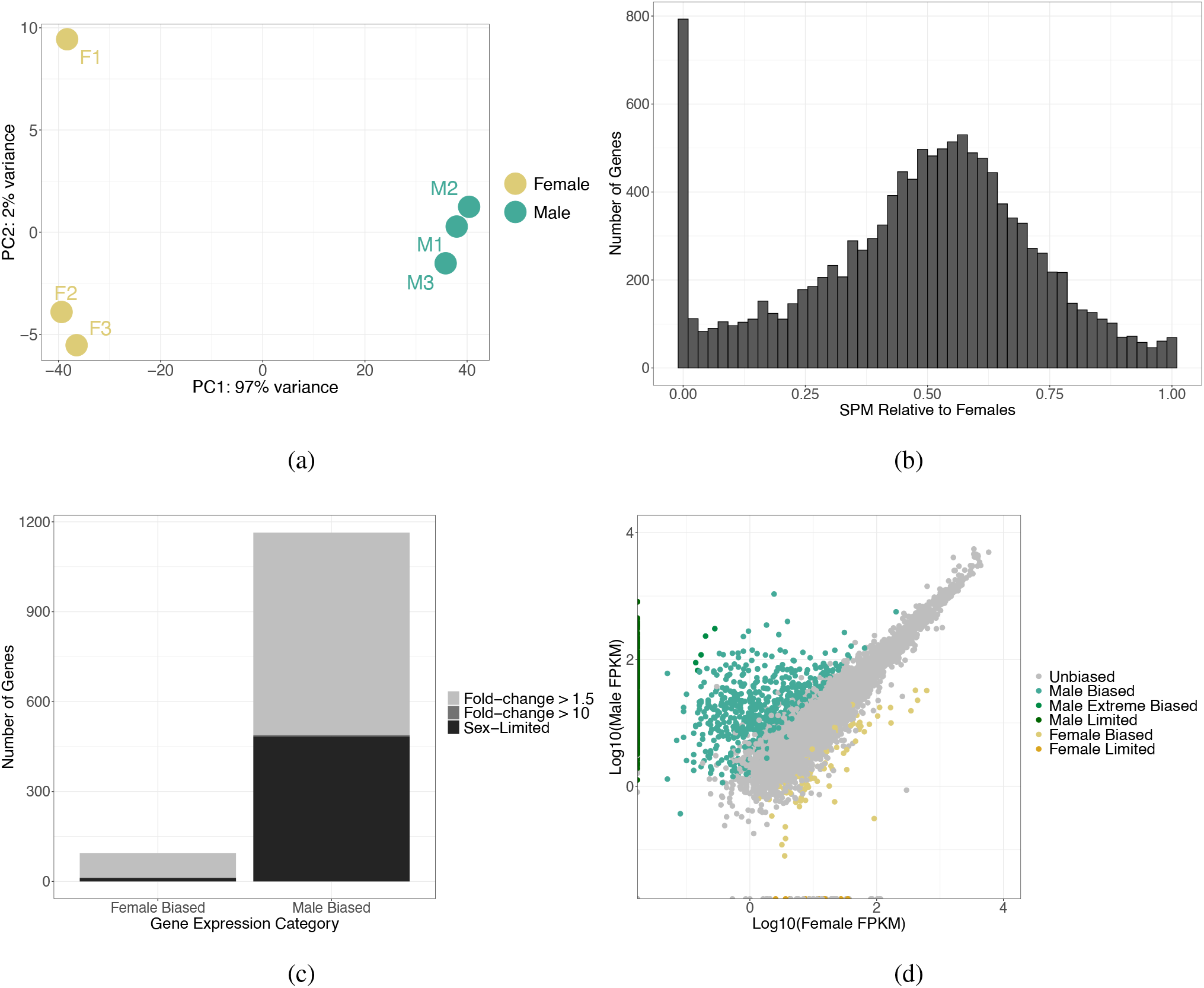
Differential gene expression between sexes. (a) PCA plot based on the expression of all genes (n = 12,420) showing 97% of the variation in expression is caused by sex. (b) Histogram of the SPM (measure of specificity (Xiao *et al*., 2010), calculated as female FPKM squared divided by female FPKM squared plus male FPKM squared) per gene (n = 12,420) showing a large number of genes are expressed only in males. (c) Stacked bar plot showing the number of sex-biased genes. Sex-limited genes referring to those with zero expression in one sex. (d) Scatter plot of the log10 fragments per kilobase of transcript per million mapped reads (FPKM) of all genes (n = 12,420). Significantly differentially expressed genes (corrected p-value <0.05 and log2 fold-change >1.5) are coloured by sex and level of differential expression, unbiased genes are shown in grey.

GO term enrichment analysis revealed differentially expressed genes from both sexes compared to all genes in the RNA-seq data set were enriched for a large variety of processes, interestingly some of these included hypermethylation of CpG islands and the regulation of various histone modifications (supplementary 1.0.2). Male-biased genes compared to all differentially expressed genes are enriched for multiple cellular processes and many regulatory processes, such as “*negative regulation of gene expression*” (GO:0010629) and “*regulation of neuron differentiation*” (GO:0045664) (supplementary 1.0.2). Female-biased genes compared to all differentially expressed genes are enriched for various biological processes and specifically reproductive related processes such as “*reproductive behaviour*” (GO:0019098) and “*pheromone biosynthetic process*” (GO:0042811) (supplementary 1.0.2).

It has been predicted that in XO sex determination systems the X chromosome may show an accumulation of male- or female-overexpressed genes which serves as a mechanism to balance the presence of sexually antagonistic alleles (Jaquiéry *et al*., 2013). We found that the X chromosome showed a depletion of male biased genes, with a significantly higher proportion of male biased genes being found on the autosomes (Chi-squared goodness of fit: X-squared = 9.171, df = 1, P-value < 0.01, Fig.3). We also found this was not the case for female biased genes, with no significant difference between the proportion of female biased genes found between the autosomes and X chromosome (Chi-squared goodness of fit: X-squared = 0.021662, df = 1, P-value = 0.88, Fig.3). These results indicate a de-masculinisation of the X chromosome in *D. citri*.

**Figure 3:**
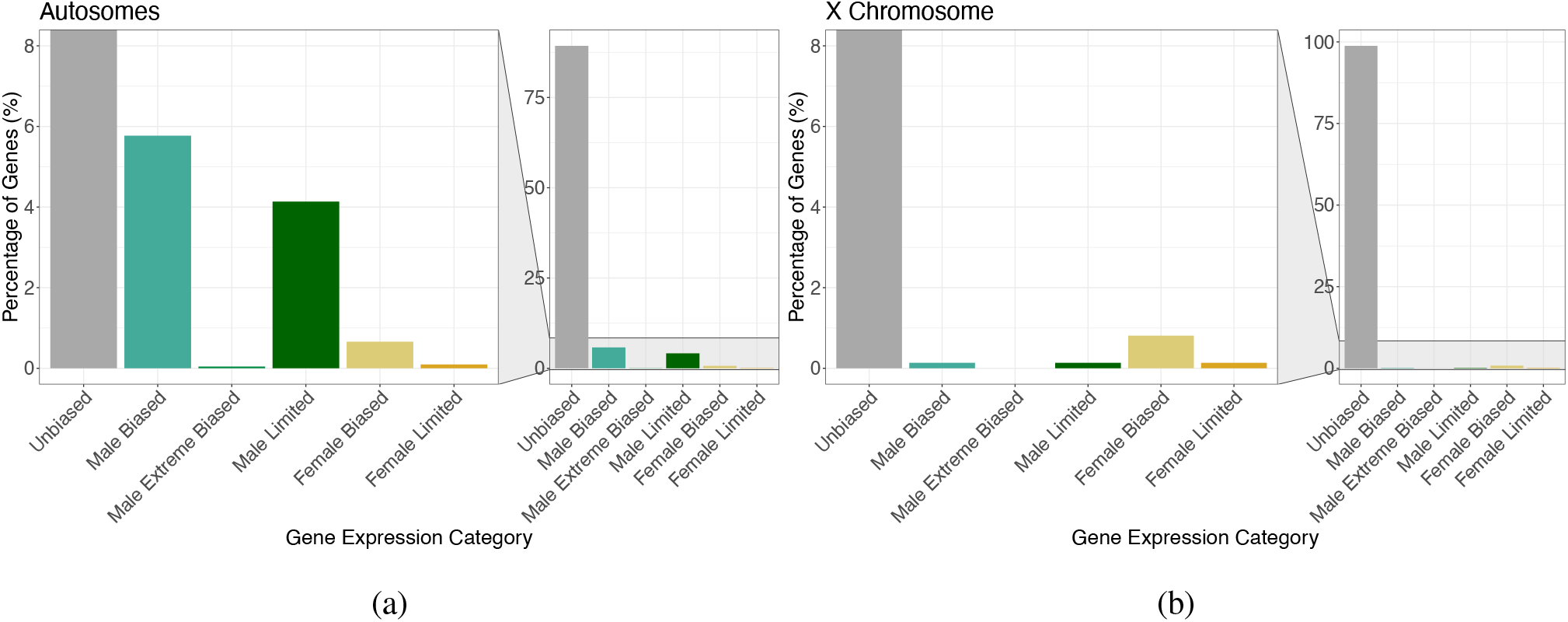
Male biased genes are depleted on the X chromosome. Bar plots showing the percentage of genes with sex-specific expression on the autosomes (a) and the X chromosome (b).

Finally, we also checked the expression levels of genes involved in DNA methylation and sexual dimorphism in *D. citri. D. citri* possesses two potential copies of *DNMT1* and no apparent *DNMT3* gene (Bewick *et al*., 2017). *DNMT1* is important for DNA methylation maintenance and *DNMT3* is involved in *de novo* DNA methylation. We blasted the *DNMT* gene sequences identified in Bewick *et al*. (2017) to the current genome annotation which resulted in two single matches, Dcitr08g10610.1.2 and Dcitr08g05090.1.1, which we will refer to as *DNMT1a* and *DNMT1b* respectively. *DNMT1b* has no detectable expression in our RNA-seq dataset for adult males or females. *DNMT1a* shows slightly higher expression in females compared to males (supplementary Fig.S3a and S3b), however overall expression is low in both sexes (<4 FPKMs) and so the difference is non-significant.

We also carried out a reciprocal blast with all *Drosophila melanogaster* isoforms of *doublesex, fruitless* and *transformer*. Whilst we find no matches for *transformer* we have identified Dcitr03g16970.1 as a *doublesex* ortholog and Dcitr01g04580.1 as a *fruitless* ortholog. There are two currently annotated isoforms for the *D. citri doublesex* ortholog which are not expressed in the adult stage of either sex. There is only one annotated isoform of the *fruitless* ortholog which is not differentially expressed between sexes (supplementary Fig.S3c) and shows overall low expression (<3 FPKMs).

### Sex-specific DNA methylation landscape of *D. citri*

Here we examine the first genome-wide methylome of a psyllid species, comparing virgin males and females. As a CpG observed/expected analysis revealed *D. citri* likely displayed DNA methylation (supplementary Fig.S4), we carried out WGBS to examine the methylome at base-pair resolution. We find low overall genome wide levels, with around 0.3% of CpGs showing methylation and zero methylation in a non-CpG context (supplementary 1.0.3 and supplementary Fig.S5). Genome-widely, males and females display similar CpG methylation profiles with some slight clustering by sex (Fig.4a). DNA methylation is also found throughout the genome in both sexes with exons showing the highest levels and intergenic regions displaying the lowest levels (Fig.4b). We specifically find a more bimodal pattern of either high or low methylation in putative promoter, UTR and exon regions compared to intergenic, intron and TE regions which show a right-skewed methylation distribution, i.e. very few regions are highly methylated (Supplementary Fig.S6 and Fig.S7).

**Figure 4:**
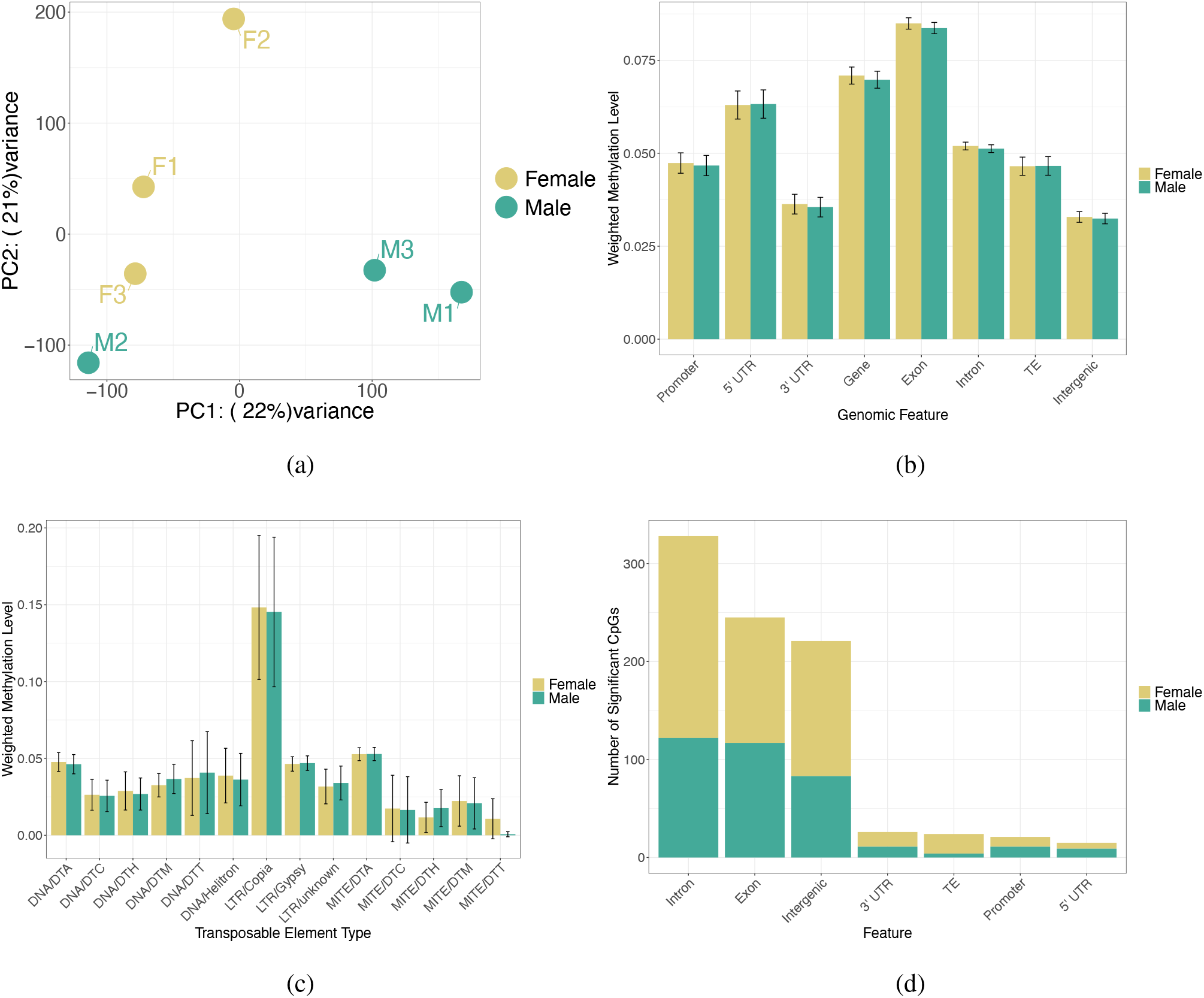
Genome-wide DNA methylation distribution in *D. citri*. (a) PCA plot based on the methylation level per CpG for all CpGs which were classed as methylated in at least one sex (n = 107,710). (b) Bar plot of the mean methylation level of each genomic feature for males and females. Error bars represent 95% confidence intervals of the mean. (c) Methylation levels of different types of TEs by sex. (d) Component bar plot showing the number of differentially methylated CpGs per genomic feature, coloured by the hypermethylated sex. Some differentially methylated CpG positions are counted twice if they overlap multiple features.

We next classified genes as showing low, medium, high and no methylation to determine if highly methylated genes show different functions to genes with lower methylation. We find highly methylated genes in both males and females are enriched for a large variety of cellular processes (supplementary 1.0.4). We also find all chromosomes, including the X, show similar proportions of genes in each methylation level category (Supplementary Fig.S8), indicating no particular chromosome is enriched or depleted for methylated genes in either males or females.

DNA methylation has been associated with transposable element silencing in other species (Zemach and Zilberman, 2010), we characterised the TE landscape of *D. citri* to examine the possibility of DNA methylation based TE regulation. We found 3.3% of the *D. citri* genome was made up of TEs with the retrotransposon *Gypsy* occupying the largest proportion of the genome, totalling around 1.5% of the autosomes and around 0.8% of the X chromosome (supplementary Fig.S9). Similar DNA methylation levels of all TEs are observed between males and females (Fig.4c), however the *Copia* class of retrotransposon shows considerably higher DNA methylation compared to all other TEs, this particular class of repeat is also only found on the autosomes and not on the X chromosome (supplementary Fig.S9).

### Sex-biased DNA methylation

A differential DNA methylation analysis of individual CpG positions between the sexes identified 763 differentially methylated CpGs (q-value <0.01 and minimum percentage difference >10%). Of these, significantly more were hypermethylated in females compared to males (Chi-squared goodness of fit: X-squared = 19.828, df = 1, p-value = <0.001.), with 443 CpGs hypermethylated in females and 320 hypermethylated in males. The majority of differentially methylated CpGs are located in genes and intergenic regions (Fig.4d). Chromosomes DC3.0sc01, DC3.0sc02 and DC3.0sc05 contain the most differentially methylated CpGs, although the number is not considerably different to all other chromosomes and there is no clear sex-bias in any specific chromosome (supplementary Fig.S10).

To create a list of confident differentially methylated features, we filtered all features to keep only those which contained at least two differentially methylated CpGs and with a minimum overall methylation difference across the entire feature of 15%. This left a final list of 12 genes containing a least one differentially methylated exon (supplementary 1.0.5). Of these 10 were hypermethylated in females and three were hypermethylated in males with one gene containing two differentially methylated exons, one hypermethylated in females and one in males (supplementary 1.0.5). None of these genes are located on the X chromosome (supplementary Fig.S11).

It is worth noting all differentially methylated genes identified above do not show overall large differences in DNA methylation (supplementary Fig.S11). Whilst we have carried out a GO term enrichment analysis for these genes, the results should be interpreted with care due to the relatively small changes in methylation. Differentially methylated genes from both sexes compared to all methylated genes are enriched for a variety of GO terms, however these terms do include, “*sex determination*” (GO:0007530), “*primary sex determination*” (GO:0007539), “*female germ-line sex determination*” (GO:0019099) and “*heterochromatin organization involved in chromatin silencing*” (GO:0070868) (supplementary 1.0.6). Genes containing hypermethylated exons in females compared to all genes containing differentially methylated exons have no enriched GO terms, and neither do genes containing hypermethylated exons in males compared to all genes containing differentially methylated exons.

### Genome-wide relationship between DNA methylation and gene expression

In many insect species, gene-body DNA methylation is positively correlated with gene expression (e.g. Bonasio *et al*., 2012; Glastad *et al*., 2016). We find this is also the case for *D. citri* with higher methylation being significantly associated with higher expression (linear model: *df* = 23971, *t* = 2.428, *p* = 0.0152, Fig.5a and 5b). The relationship between gene expression and methylation is similar in both sexes as there is no significant interaction between sex and methylation level (two-way ANOVA: *F_2, 23971_* = 2.952, *p* = 0.433). On a genome-wide scale, it is clear that this relationship is conserved in only the most highly methylated genes (Fig.5c and 5d).

**Figure 5:**
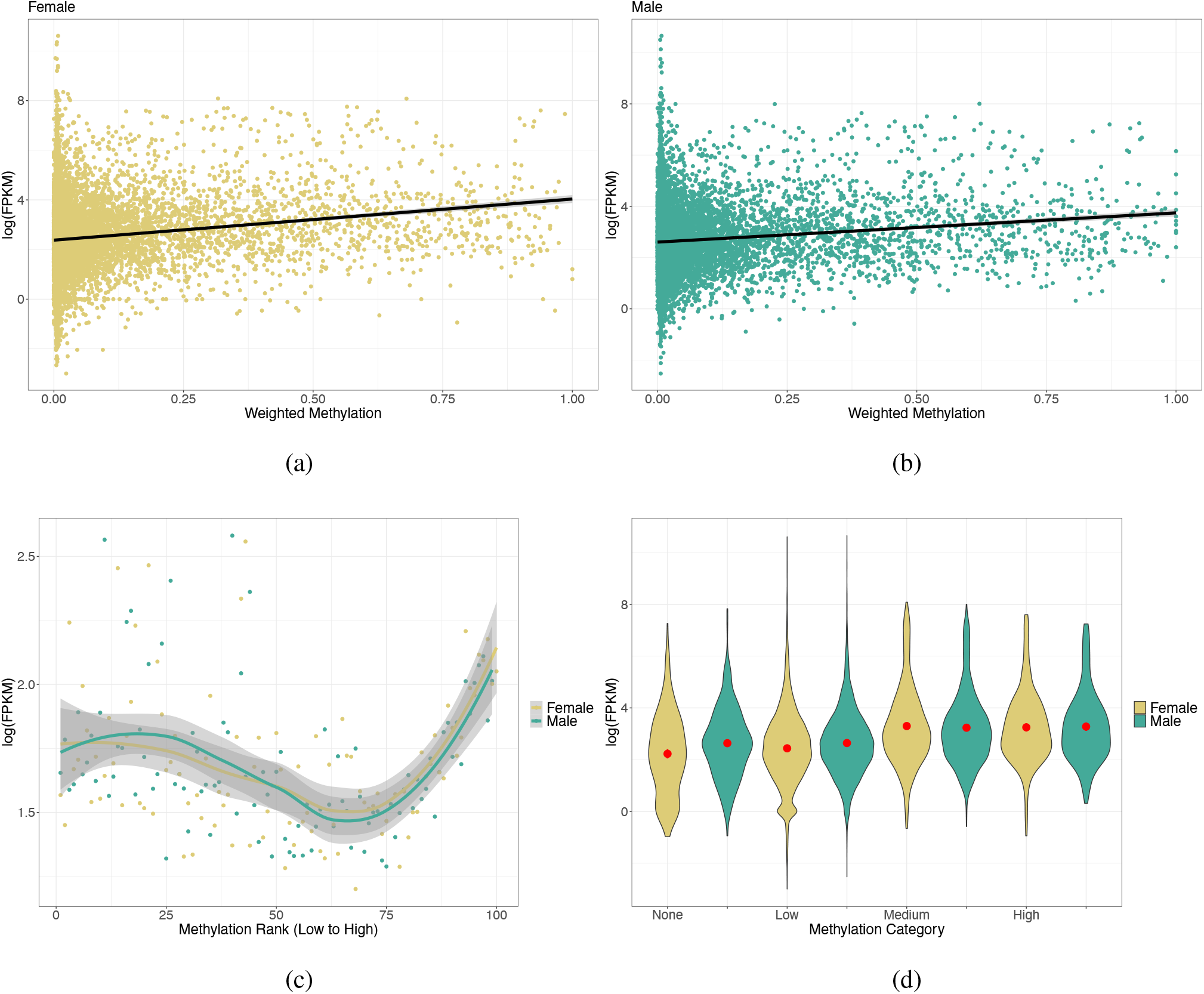
Genome-wide DNA methylation and gene expression relationship. (a and b) Scatter graphs of the mean weighted methylation level per gene (averaged across replicates) plotted against the mean expression level. Each dot represents a gene, the black lines show a fitted linear regression with grey areas indicating 95% confidence intervals. (c) Binned genes by mean weighted methylation level with the mean expression level plotted for each bin with fitted LOESS regression lines per sex. Grey areas indicate 95% confidence areas. (d) Violin plots showing the distribution of the data via a mirrored density plot, meaning the widest part of the plots represent the most genes. Weighted methylation level per gene per sex, averaged across replicates, was binned into four categories, no methylation, low (>0-0.3), medium (>0.3-0.7) and high (>0.7-1). The red dot indicates the mean with 95% confidence intervals.

We also examined the relationship between DNA methylation and expression separately for the autosomes and the X chromosome. We find that the association between methylation and expression is only significant for the autosomes, and not the genes on the X chromosome (Autosomes: linear model: *df* = 22543, *t* = 2.538, *p* = 0.0112, X chromosomes: linear model: *df* = 1425, *t* = −1.692, *p* = 0.0909, supplementary 2: Fig.S12 and S13.).

### Relationship between sex-specific DNA methylation and expression

The role of differential DNA methylation in regulating differential gene expression between insect sexes appears to differ between species (Mathers *et al*., 2019; Bain *et al*., 2021). We therefore searched for a potential relationship between differential exon DNA methylation and sex-specific gene expression in *D. citri*. We find no difference in the expression levels of genes which are differentially methylated or not (linear model: *df* = 23958, *t* = 0.183, *p* = 0.99, Fig.6a). We do, however, find genes with unbiased expression have significantly higher levels of DNA methylation compared to differentially expressed genes (linear model: *df* = 23968, *t* = 3.893, *p* < 0.01, Fig.6b), this effect is not sex-specific (two-way ANOVA: *F_2,23968_* = 0.0122, *p* = 0.9879). Finally, on a single gene level there is no correlation between differential DNA methylation and differential gene expression between the sexes (supplementary Fig.S14).

**Figure 6:**
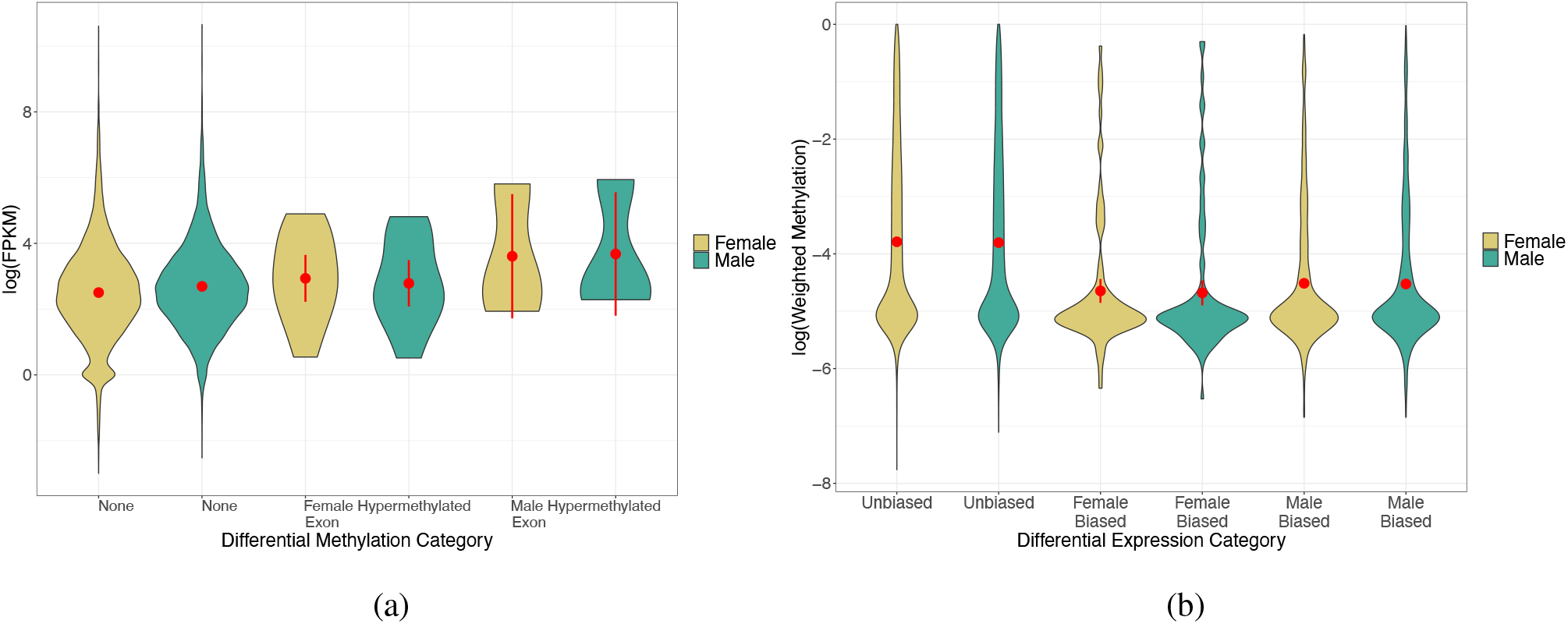
Relationship between differential DNA methylation and differential expression. (a) Violin plot of the expression levels of genes which are differentially methylated or not between sexes. The red dot represents the mean with 95% confidence intervals. (b) Violin plot of the methylation levels of genes which are either unbiased or show sex-specific expression bias. The red dot represents the mean with 95% confidence intervals.

## Discussion

In this study, we present the first detailed analysis of genome-wide sex-specific DNA methylation patterns in the agriculturally important insect pest, *D. citri*, evaluating its effects on gene expression and sexual differentiation. Our major findings include: the identification of the X chromosome (chromosome 08 in Diaci_v3), sex-biased gene expression characterized by a large number of male limited genes, a depletion of male-biased genes on the X chromosome, overall low genome-wide levels of DNA methylation, with DNA methylation targeted to exons but also present in promoters and TEs, a small number of differentially methylated genes between the sexes and no apparent *cis*-driven relationship between differential DNA methylation and differential gene expression.

### Sex-biased gene expression in an X0 system

*D. citri* harbors an XX/X0 sex determination system, whereby females carry two X chromosomes and males only one X. Aside from initial sex determination, genes on the sex chromosomes are theorized to play a disproportionately large role in phenotypic differences between males and females (Dean and Mank, 2014). Here, we found a de-masculinisation of the X chromosome in *D. citri* indicated by a reduction in genes showing male biased expression relative to the autosomes. A similar de-masculinisation of the X has been observed in many *Drosophila* species (Sturgill *et al*., 2007), nematodes (Reinke *et al*., 2004), as well as in Hemiptera, *Halyomorpha halys* and *Oncopeltus fasciatus* (Pal and Vicoso, 2015). This is in line with classic evolutionary theories that hold the X chromosome, whose sex-biased transmission sees it spending more time in females, should value females more than males by de-masculinisation and/or feminization (Hitchcock and Gardner, 2020). However, the widespread application of next-generation sequencing (NGS) techniques on different organisms has revealed inconsistent patterns among species. We did not find any evidence for a feminized X chromosome, based on gene-expression differences, which was also predicted by these theories. Empirical evidences demonstrated that female-biased genes were under-represented on the X chromosomes of nematodes and a masculinized X was found in the pea aphid *Acyrthosiphon pisum*, directly contradicting evolutionary theories (Jaquiéry *et al*., 2013; Reinke *et al*., 2004). Considering that a gene’s impact on phenotype may become diluted when it moves from a haploid to a diploid setting (Otto, 2007), recent theoretical predictions suggest the relative power of an X-linked gene to induce fitness effects may be lower in a female carrier than in a male. This power asymmetry is proposed to generate a bias towards male-beneficial strategies that offset, and even overturn, the X-linked gene’s feminization (Jaquiéry *et al*., 2013; Hitchcock and Gardner, 2020), which may explain the interesting observations regarding female-biased genes on the X chromosome were not overrepresented in this study and even enrichment of male-biased genes on X in *A. pisum*. Noteworthily, the whole-body comparisons of RNA-seq data between sexes are somewhat limited, and potentially introduce biases in the analysis (Pal and Vicoso, 2015; Perry *et al*., 2014). Further comparisons using the data from different male and female tissues will be necessary to confirm the extent of de-masculinisation on X chromosome in *D. citri*.

### Similar DNA methylation profiles between males and females

In addition to characterising chromosome-specific differential expression between sexes, we also looked at sex-specific genome-wide DNA methylation differences. We find overall considerably lower levels of DNA methylation in *D. citri* compared to other hemipteran insects which generally have been found to show >2% CpG methylation (Bewick *et al*., 2017; Mathers *et al*., 2019; Bain *et al*., 2021). The low levels found here (0.3%) more closely match the low levels found in Hymenoptera and Lepidoptera (Bewick *et al*., 2017). This shows the importance of investigating epigenetic profiles in individual species and not making assumptions based on related species. This idea is also highlighted by the recent finding of promoter DNA methylation in some insects (Lewis *et al*., 2020) for which we also see some evidence for in *D. citri*, although it should be noted that the levels are similar to background intergenic levels.

We also find no difference in DNA methylation profiles between the autosomes and the X chromosome, this has rarely been investigated to date due to the lack of chromosome level assemblies for non-model insects. However, Mathers *et al*. (2019) do find a depletion in highly methylated genes on the X chromosome of a species of aphid, again these differences highlight the diversity of species-specific epigenetic profiles. Additionally, we find no genome-wide sex differences across genomic regions between sexes. This is similar to two jewel wasp species, *Nasonia vitripennis* and *Nasonia giraulti*, in which more than 75% of expressed genes displayed sex-biased expression, but no sex differences in DNA methylation were observed (Wang *et al*., 2015). However, extreme sex-biased DNA methylation has been observed in many insect systems including *M. persicae, Zootermopsis nevadensis* and *P. citri* (Mathers *et al*., 2019; Glastad *et al*., 2016; Bain *et al*., 2021) Examples include a unique sex-specific pattern in *P. citri*, in which males display more uniform low levels of methylation across the genome, while females display more targeted high levels (Bain *et al*., 2021). The fact that DNA methylation is so similar between the sexes may be considered when developing a molecular control system, e.g. RNAi strategy against *D. citri* as a pest species. RNAi-mediated gene knockdown has shown tremendous potential for controlling the hemipteran aphids and psyllids (Jain *et al*., 2021; Yu and Killiny, 2020; Yu *et al*., 2016), our results may indicate that RNAi of a target gene should have similar silencing efficiency in both sexes of *D. citri*. It has also recently been shown that knockdown of *DNMT1* in *Phenacoccus solenopsis* by RNAi resulted in offspring death (Omar *et al*., 2020); similar observations were recorded in another hemipteran insect, *Nilaparvata lugens*, that silencing *DNMT1* and *DNMT3* caused fewer offspring (Zhang *et al*., 2015), suggesting such a strategy may hold potential for the control of *D. citri*.

### Conserved TE methylation

In order to explore TE methylation, we characterised TEs with the *D. citri* genome. An interesting outcome of our study is the low TE proportion in *D. citri*. TEs, known as jumping genes propagating in genomes, are associated with a variety of mechanisms contributing to shape genome architecture and evolution (Gilbert *et al*., 2021). In insects, TEs mediate genomic changes which have been reported to play a pivotal role in the development of insecticide resistance, as well as adaptation to climate change, and local adaptation (Adrion *et al*., 2019; González *et al*., 2010; Itokawa *et al*., 2010). Dedicated comparative analyses of TE composition reveals insect TE landscapes are highly variable between insect orders and among species of the same order. The genomic portion of TEs ranges from as little as 0.12% in the antarctic midge, *Belgica antarctica*, to as large as 60% in the migratory locust *Locusta migratoria* (Kelley *et al*., 2014; Petersen *et al*., 2019; Wang *et al*., 2014). Even within closely related species, TE composition can be drastically different; *Aedes aegypti* TEs contribute about 47% of the whole genome, followed by 29% in *Culex quinquefasciatus*, 20% in *D. melanogaster*, 16% in *Anopheles gambiae* and 0.12% in *B. Antarctica* (Arensburger *et al*., 2010; Kelley *et al*., 2014; Nene *et al*., 2007; Quesneville *et al*., 2005; Sharakhova *et al*., 2007). In this study, we found only 3.3% of the *D. citri* genome was made up of TEs, a small proportion compared with other reported hemipteran insects such as *Cimex lectularius* (30%), *A. pisum* (25%), *H. halys* (39%), *Pachypsylla venusta* (24%) and *O.fasciatus* (21%) (Petersen *et al*., 2019). By investigating the 195 insect genomes, a study uncovered large-scale horizontal transfer of TEs from host plants or a bacterial/viral infection (Peccoud *et al*., 2017), and makes this mechanism likely to be the source of high variation in insect genomic TE composition. Meanwhile, TE content is usually positively correlated with arthropod genome size (Gilbert *et al*., 2021; Petersen *et al*., 2019); and *D. citri* does indeed show a relatively small genome, at around 475Mb (Hosmani *et al*., 2019).

We additionally find evidence of TE methylation in *D. citri*. TE methylation is generally found across plants and animals (Law and Jacobsen, 2010), however, it was thought to be lost in arthropods (Keller *et al*., 2016; Zemach *et al*., 2010), although this was based on a small number of investigated species. Recently, TE methylation has been shown to be present in centipedes and a species of mealybug (Lewis *et al*., 2020) as well as in the desert locust (Falckenhayn *et al*., 2013). Whilst, we also show TE methylation in *D. citri*. It is worth remembering the genome-wide level of DNA methylation in *D. citri* is particularly low (0.3%) and as such methylation of TEs may not be functioning to silence TE movement as in other highly methylated species. This discovery does however add to the growing evidence that the function of DNA methylation is highly variable between insect species.

### DNA methylation does not drive sex-specific gene expression

Finally, we have identified the relationship between DNA methylation and gene expression in *D. citri*. We find highly methylated genes show generally higher levels of gene expression, which appears common within insects (e.g. Bonasio *et al*., 2012; Glastad *et al*., 2016; Marshall *et al*., 2019), although see Bain *et al*. (2021). This trend is common between both sexes and we also find no relationship between differential DNA methylation and differential gene expression, again this has been shown to be the case in multiple other insect studies exploring sex-specific DNA methylation profiles (Wang *et al*., 2015; Glastad *et al*., 2016), although see Mathers *et al*. (2019).

It has recently been suggested that DNA methylation may play a temporal role in regulating gene expression (Li-Byarlay *et al*., 2020), whereby DNA methylation creates changes in chromatin structure through the recruitment of histone modifications (Xu *et al*., 2021) and this allows a later change in gene expression which would not be present in samples taken during the same time frame. The initial DNA methylation event may then be lost accounting for the lack of association between gene expression and DNA methylation, although this idea is yet to be tested. If DNA methylation were functioning in this temporal fashion in *D. citri*, we may expect to see DNA methylation differences between different developmental stages. Indeed, the small number of differentially methylated genes we have identified between sexes are involved in sex differentiation and heterochromatin formation. As we used adult whole-bodies in this study the DNA methylation differences in these genes may be due to tissue-specific profiles (Pai *et al*., 2011). Although, there is growing evidence that the underlying genomic sequence drives DNA methylation patterns in many insect species (Yagound *et al*., 2020; Harris *et al*., 2019; Marshall *et al*., 2019; Yagound *et al*., 2019). Future work sampling different tissues and developmental stages may shed light on a potential role for DNA methylation in earlier development, which would allow for a more targeted approach to epigenetic pest control.

## Conclusion

This study provides a fundamental basis for future research exploring epigenetic mechanisms of insect control in an important agricultural pest species, *D. citri*. We have further characterised the current *D. citri* reference genome by identifying the X chromosome in this species and explored the TE content, finding low genome-wide TE levels. We also find a large number of genes show male-biased expression and find the X-chromosome is depleted for male-biased genes. Importantly, we characterise the sex-specific methylome of *D. citri* finding evidence for promoter and TE methylation, although genome-wide *D. citri* shows considerably lower DNA methylation levels than most of hemipteran species currently studied. Given that the small number of differentially methylated genes we do find between sexes are involved in processes such as sex differentiation, we suggest DNA methylation may play a more functional role in earlier developmental stages. Finally, we find no relationship between *cis*-acting DNA methylation and differential gene expression, as is common in many insects. The similar DNA methylation profiles between sexes reported here would help to develop an epigenetic-based pest control method targeted at DNA methylation for *D. citri* management.

## Acknowledgements

We thank Dr. Kamil Jaron for valuable advice regarding X chromosome identification. This work is funded by the National Natural Science Foundation of China (grant no. 32160634 and 31601379), the Natural Science Foundation for Distinguished Young Scholars of Jiangxi province (grant no. 20212ACB215001), and the Major Science and Technology R&D Program of Jiangxi Province (grant no. 20194ABC28007). HM was funded by the ERC Grant ‘PGRrepro’ awarded to LR.

## Author contributions

XY, HM, LR and ZL conceived the study. XY, YL, YX, XZ, HY, WC, GZ and BZ reared the insects and conducted the laboratory work. HM carried out all analyses. XY and HM wrote the initial manuscript. All authors contributed to the final manuscript.

## Data Accessibility

All RNA-seq, BS-seq and whole genome sequencing data generated for this study were deposited in GenBank under Bio-Project accession number PRJNA774108. All scripts are available at: https://github.com/MooHoll/Asian_Psyllid_Methylation. New genome annotations and GO terms can be found at: https://doi.org/10.6084/m9.figshare.17013980.v1.

