## Supplementary_2 for "Sex-specific transcription and DNA methylation landscapes of the Asian citrus psyllid, a vector of huanglongbing pathogens"

### Supplementary 2.0: additional figures

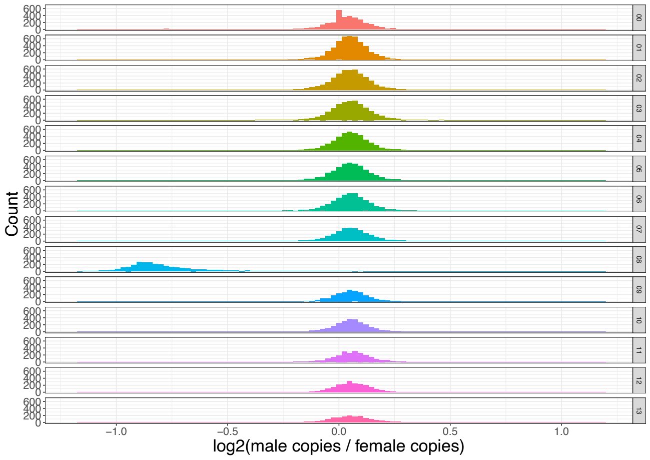


(a)

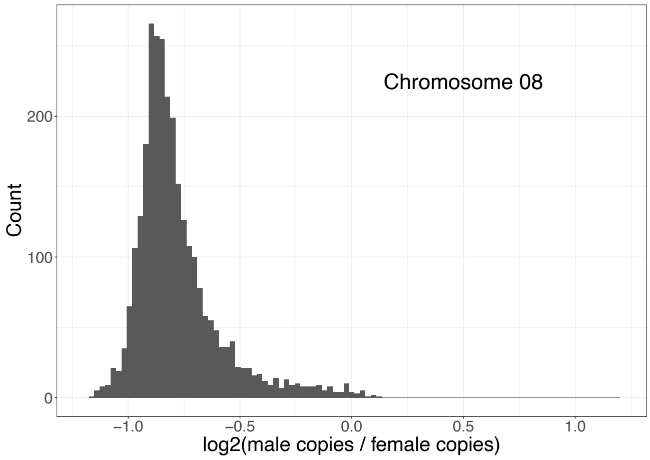


(b)

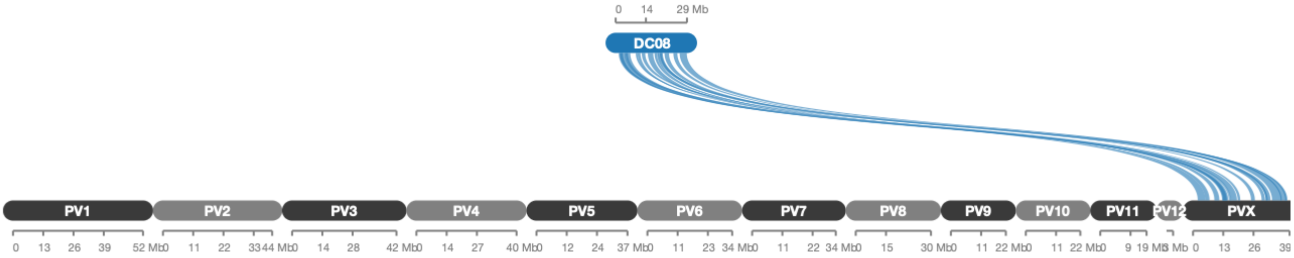


(c)

Figure S1: **Identification of the X chromosome.** (a) Histogram of the log2 male to female coverage ratio for 10,000bp windows across each chromosome. (b) Histogram of the log2 male to female coverage ratio for 10,000bp windows for chromosome 08 only. (c) Synteny plot showing collinearity blocks between *D. citri* (DC) chromosome 08 and *P. venusta* (PV). Each line represents at least 10 orthologous genes.

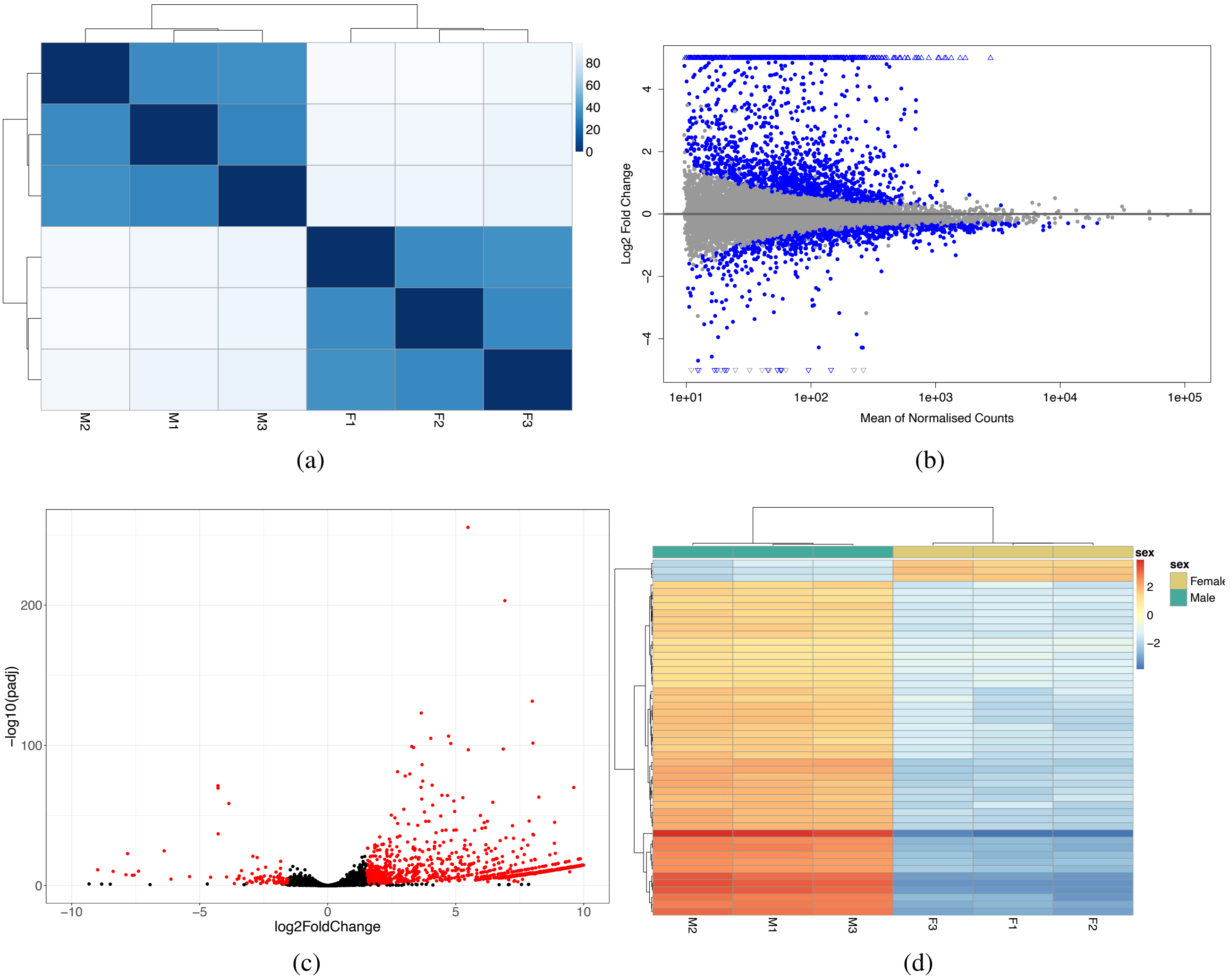


Figure S2: **Differential gene expression between sexes.** (a) Poisson measurement of dissimilarity between counts for all samples. (b) MA plot showing differentially expressed genes highlighted in blue. (c) Volcano plot showing differentially expressed genes shown in red. (d) Heatmap showing the top 50 differentially expressed genes between sexes.

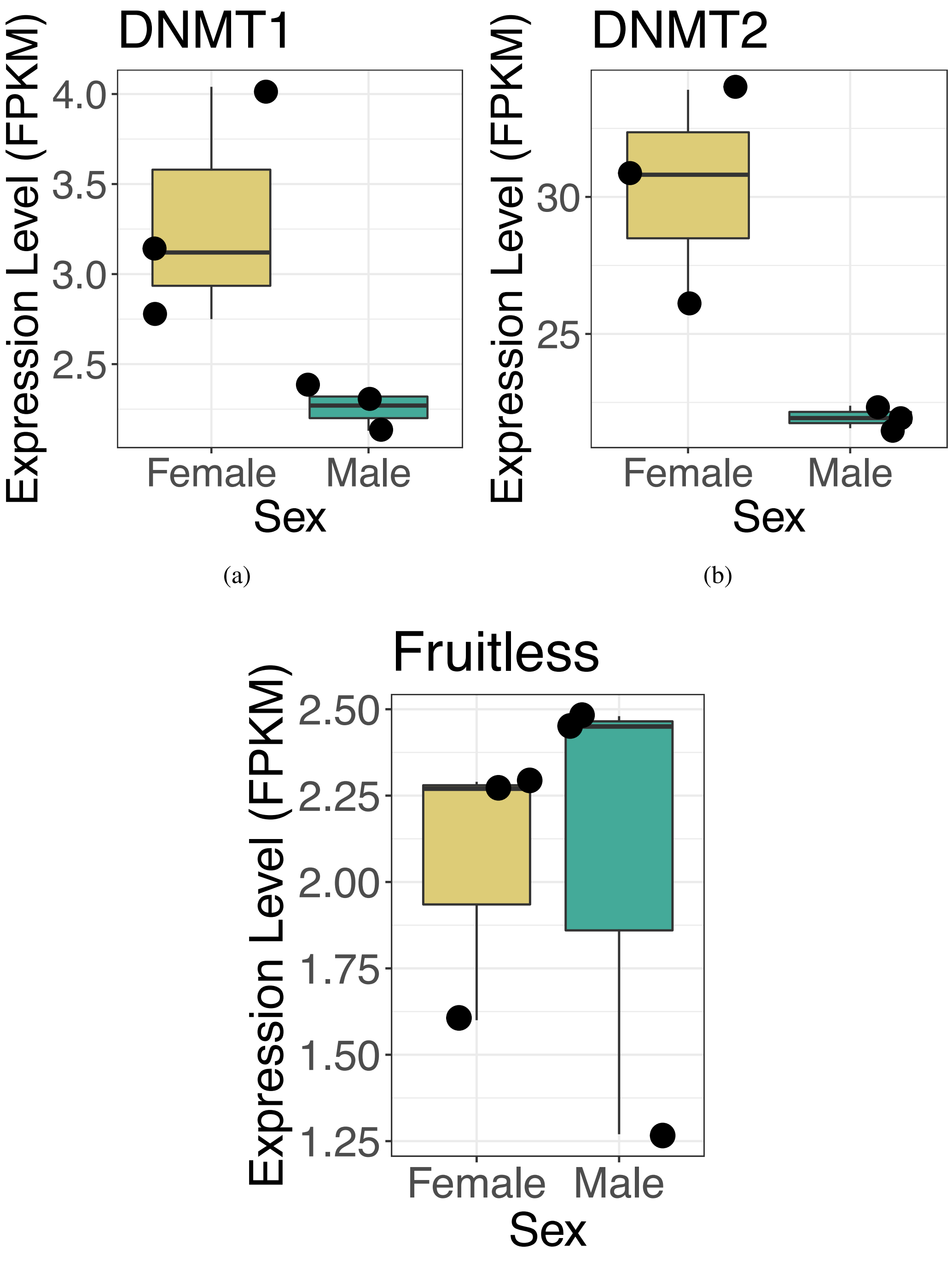


(a) (b)

(c)

Figure S3: **Expression of specific genes.** Boxplots showing the expression levels of the *DNMT1a* (a), *DNMT2* (b) and *fruitless* (c) genes. *DNMT1b* is not plotted as there was no detectable expression in either sex. Whilst both DNMT genes are more highly expressed in females the difference is non-significant (adjusted p-value >0.05). It is also worth nothing *DNMT2* is usually associated with tRNA methylation not DNA methylation.

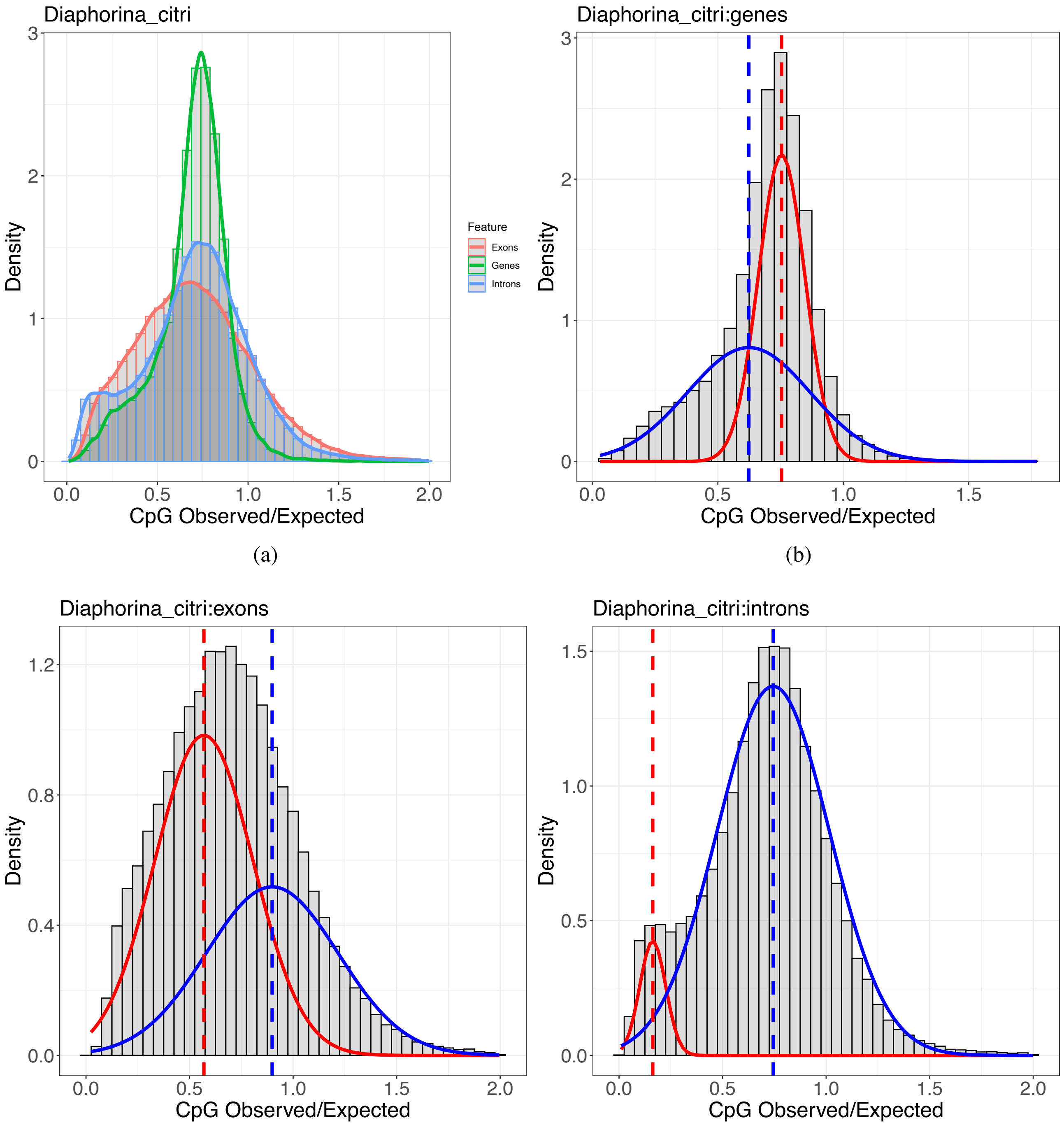


(a) (b) (c) (d)

Figure S4: **CpG observed/expected (o/e) density graphs.** a) the density of CpGs present in a region give the CG content of that region, for genes, exons and introns. A peak at 1.0 indicates likely no DNA methylation is present a peak at 0.5 indicates the presence of DNA methylation. A decrease in expected CpG positions is caused by an increase in deamination of cytosine bases to thymine bases when DNA methylation is present. b) CpG o/e for genes, c) exons and d) introns with two distributions identified by the R package *mixtools* v1.2.0.

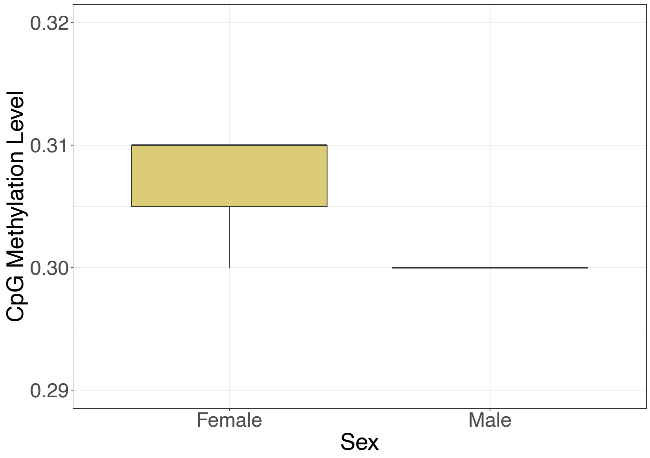


Figure S5: **Genome-wide methylation patterns.** Boxplots showing the mean genome-wide levels of CpG DNA methylation per biological replicate per sex. Calculated as the proportion of reads supporting a cytosine call per CpG position.

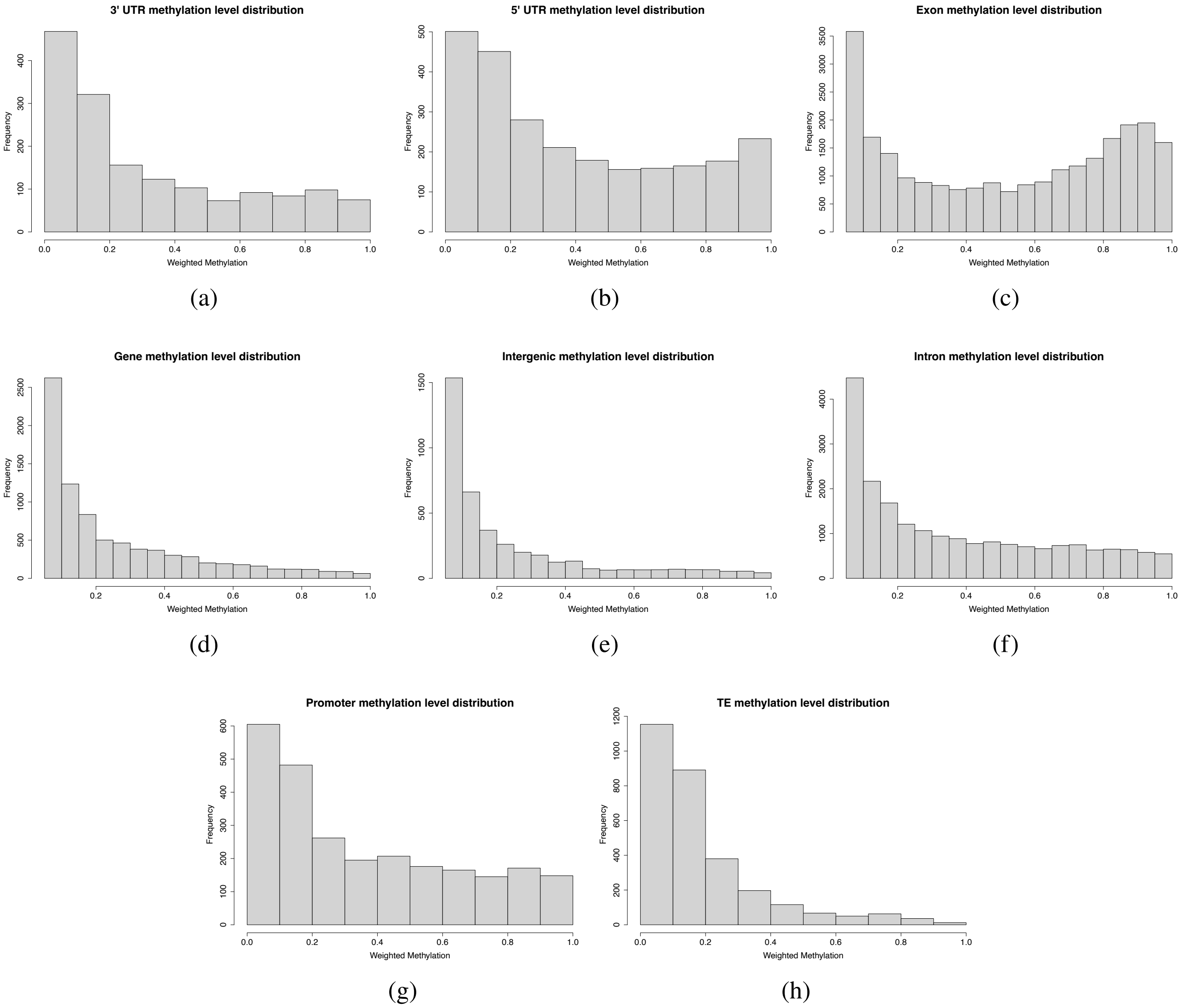


Figure S6: **Distribution of methylation levels across methylated features.** Histograms showing the distribution of methylation levels for genomic features, (a) 3' UTRs, (b) 5' UTRs, (c) exons, (d) whole genes, (e) intergenic regions, (f) introns, (g) putative promoters and (h) TEs. All graphs only show features with a weighted methylation level greater than the lambda conversion rate of 0.05.

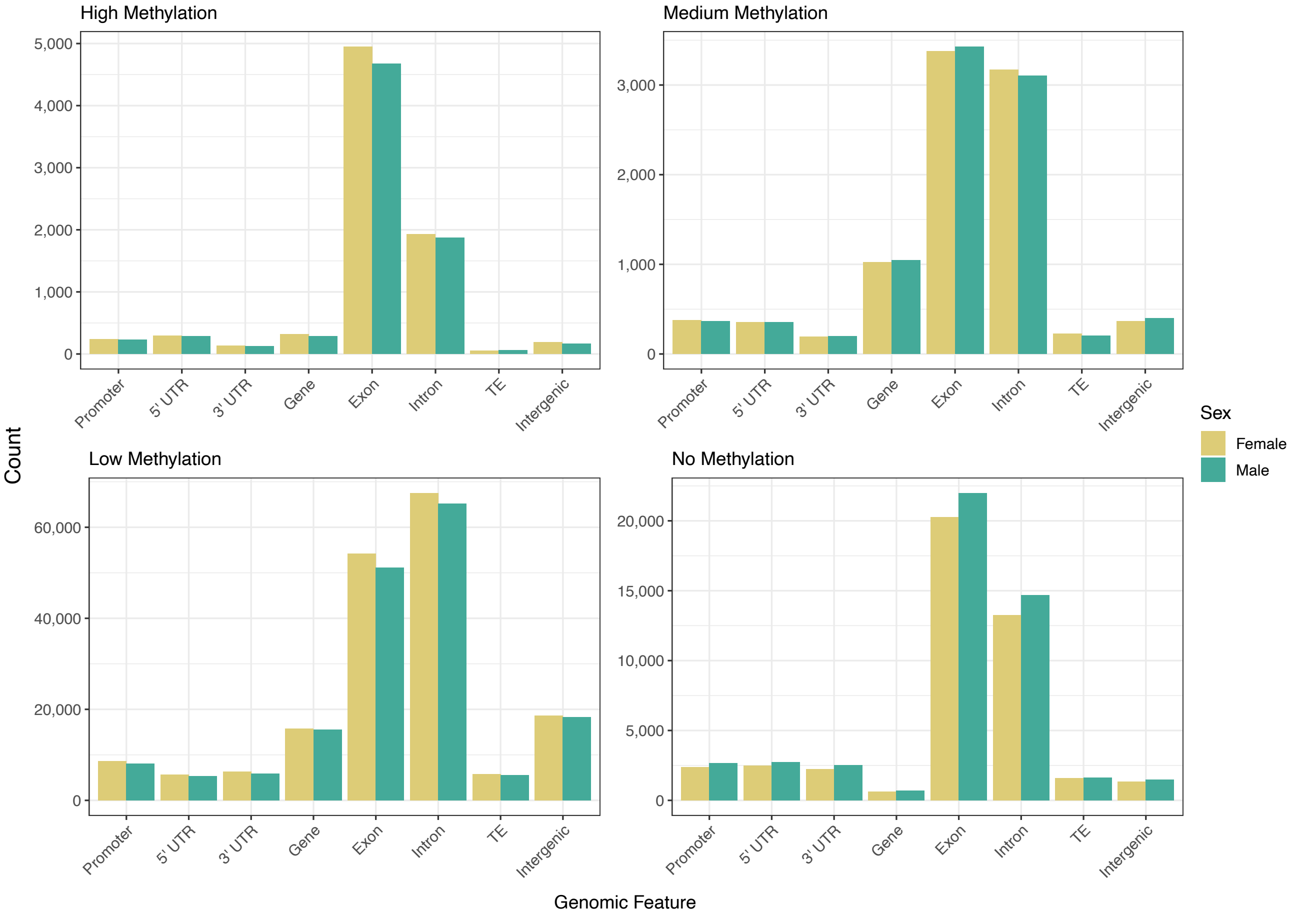


Figure S7: **Counts of features with different levels of methylation.** Bar plots of the total number of genomic features categorised by weighted methylation level for males and females. High methylation is a weighted methylation level >0.7, medium is >0.3-0.7, low is >0-0.3 and no methylation is equal to zero.

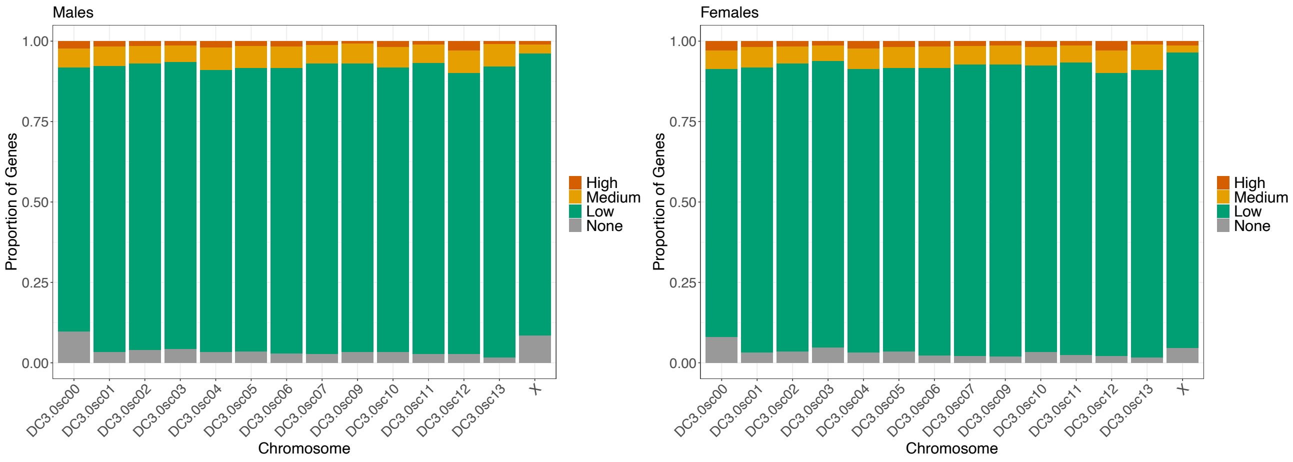


(a) (b)

Figure S8: **Proportion of methylated genes per chromosome based on exon methylation levels.** Stacked bar chart showing the proportion of genes per chromosome which have different levels of exon methylation for (a) males and (b) females. A methylation level of 'None' refers to zero methylation present, low is a weighted methylation level of >0-0.3, medium is >0.3-0.7 and high is >0.7.

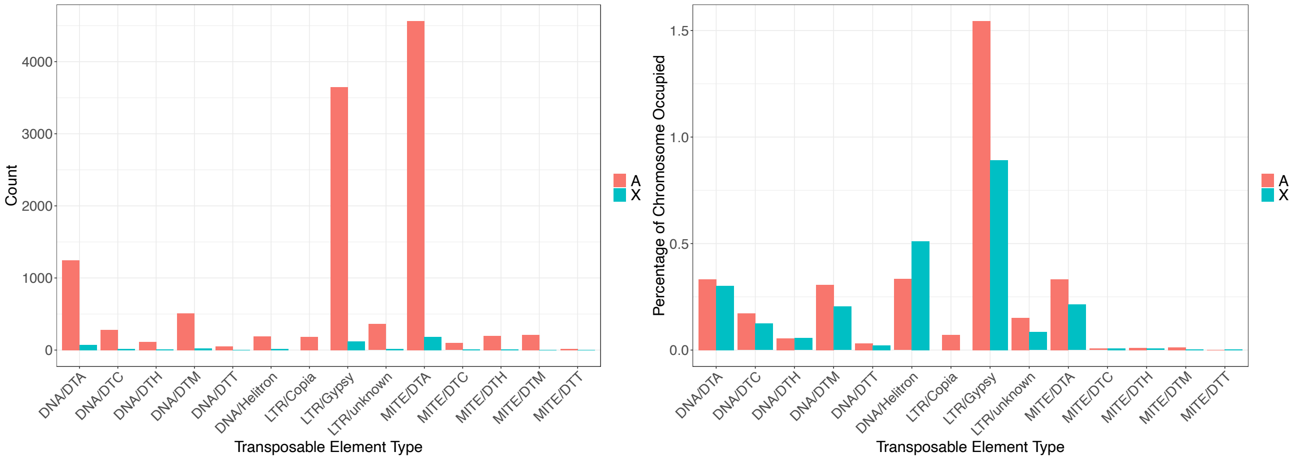


(a) (b)

Figure S9: **TE distribution and methylation levels.** (a) Total number of TEs by category found within autosomes and the X chromosome. (b) percentage of the chromosomes (either autosomes or X chromosome) occupied by TEs.

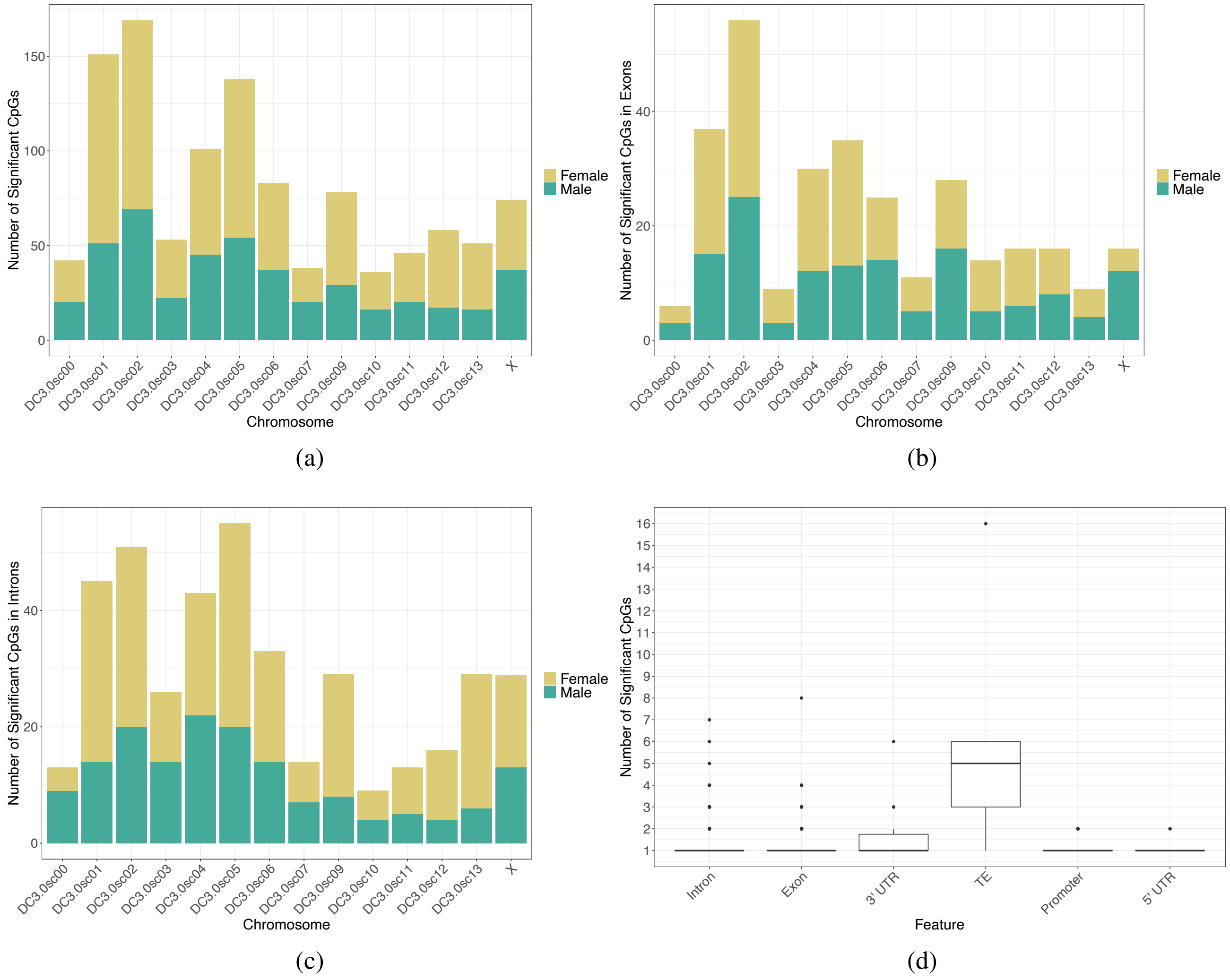


Figure S10: **Location of differentially methylated CpGs by chromosome.** (a) location of all differentially methylated CpGs by chromosome, hypermethylated CpGs are coloured by sex. (b) CpGs in exons only. (c) CpGs in introns only. (d) Boxplot of the number of differentially methylated CpGs per feature per gene.

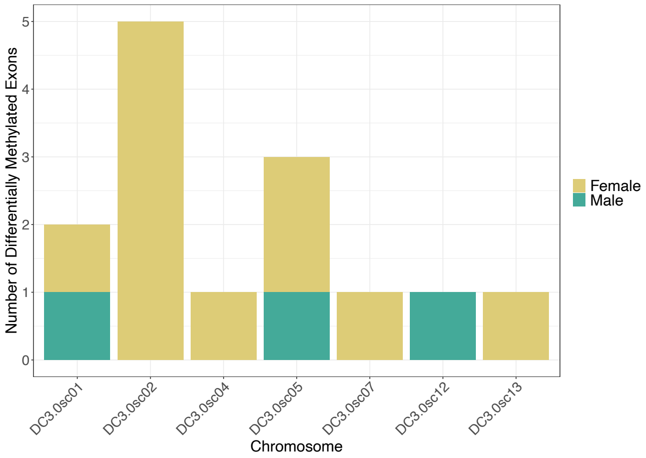


(a)

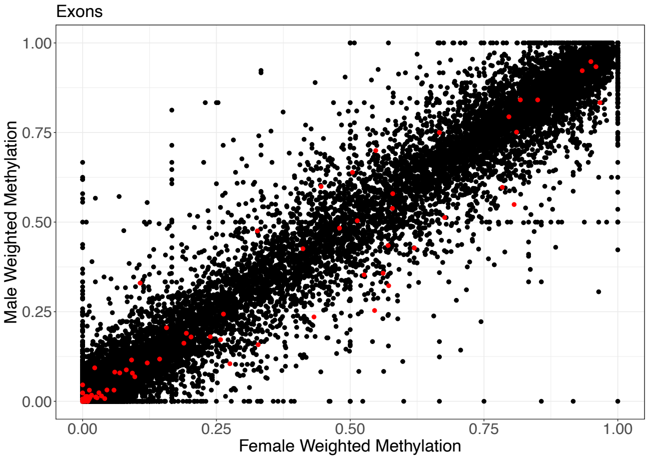


(b)

Figure S11: **Differentially methylated exons.** (a) Chromosome location of genes with differentially methylated exons. (b) Scatter plot of the mean weighted methylation levels of exons for males and females. Red dots indicate significant differentially methylated features (containing two significant differentially methylated CpGs and a minimum weighted methylation difference of 15%)

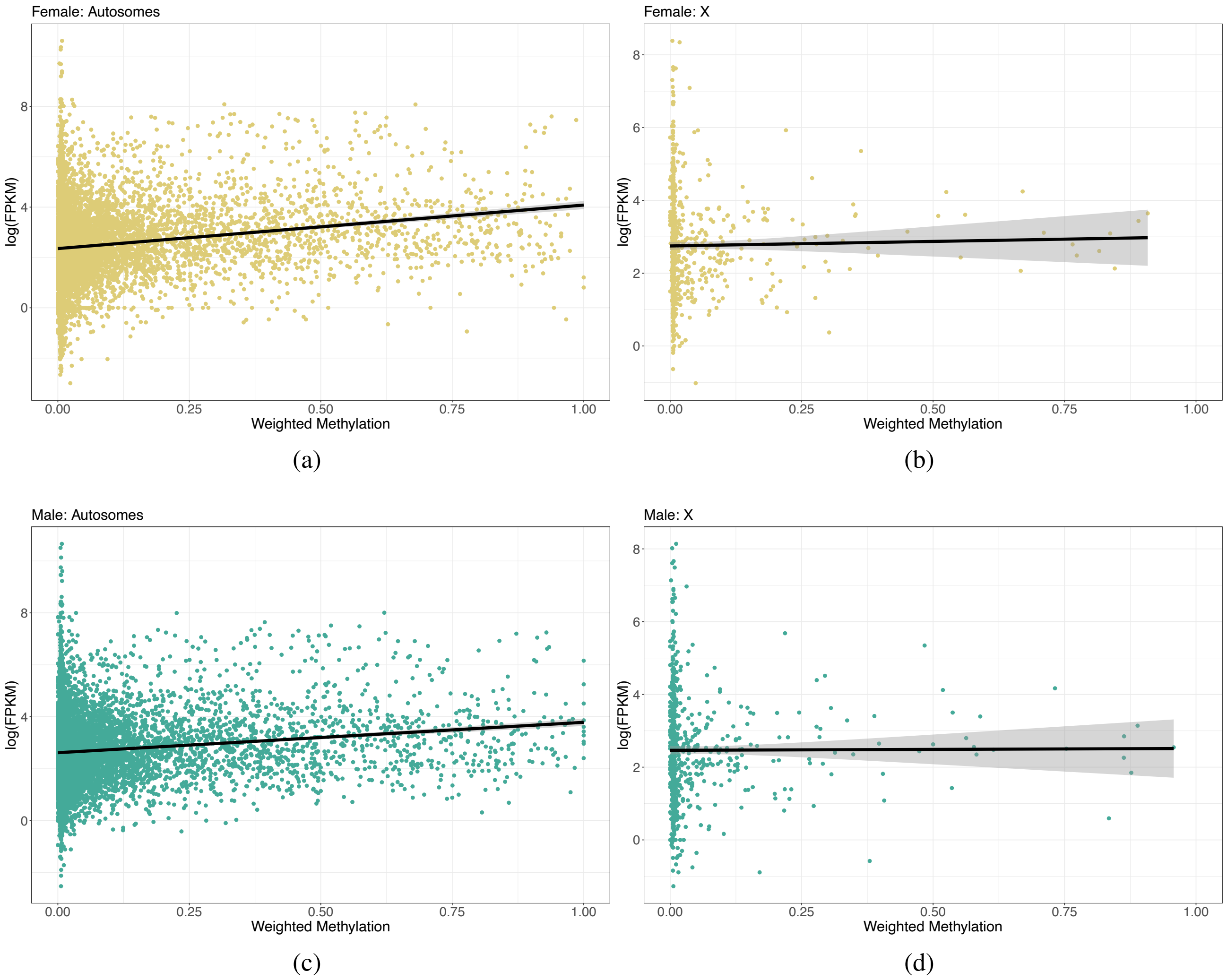


Figure S12: **Genome-wide relationship between gene expression and DNA methylation for both sexes separated by autosomes and the X chromosome.** Scatter graphs of the mean weighted methylation level per gene (averaged across replicates) plotted against the mean expression level, (a) female autosomes, (b) female X chromosome, (c) male autosomes and (d) male X chromosome. Each dot represents a genes, the black lines show a fitted linear regression with grey areas indicating 95% confidence intervals.

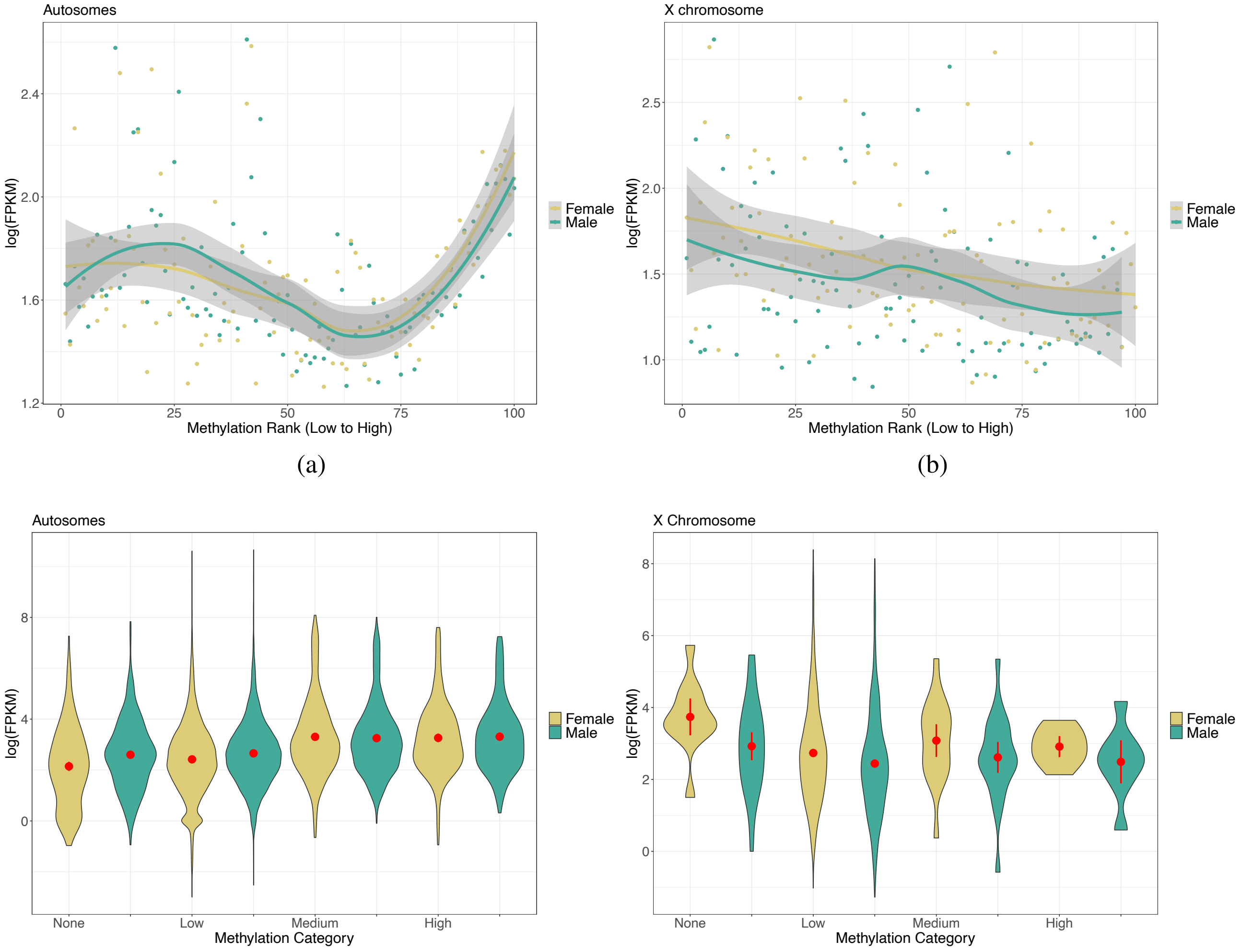


(a) (b)

(c) (d)

Figure S13: **Genome-wide relationship between gene expression and DNA methylation for both sexes separated by autosomes and the X chromosome.** (a and b) Binned genes by mean weighted methylation level with the mean expression level plotted for each bin with fitted LOESS regression lines per sex. Grey areas indicate 95% confidence areas. (c and d) Violin plots showing the distribution of the data via a mirrored density plot, meaning the widest part of the plots represent the most genes. Weighted methylation level per promoter per sex, averaged across replicates, was binned into four categories, no methylation, low (>0–0.3), medium (>0.3–0.7) and high (>0.7–1). The red dot indicates the mean with 95% confidence intervals.

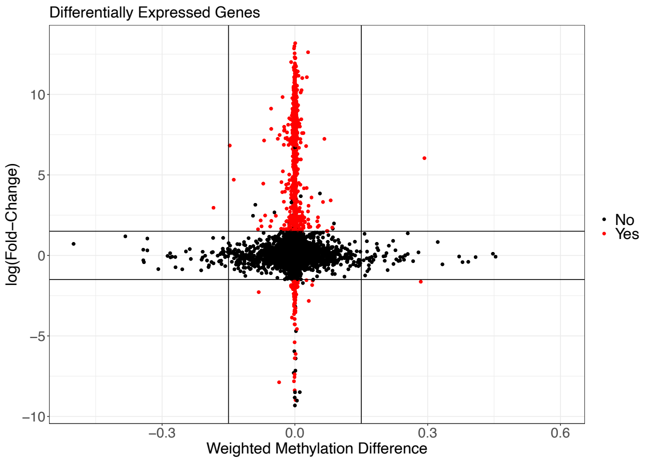


(a)

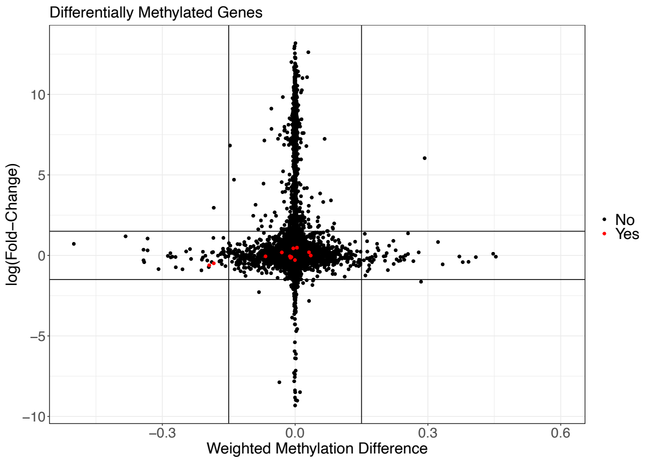


(b)

Figure S14: **Methylation and gene expression differences between *D. citri* sexes.** Scatter plot of the weighted methylation difference between sexes (mean male weighted methylation minus mean female weighted methylation) for genes plotted against the log fold-change in gene expression. A log fold-change greater than zero represents overexpression in males. Each point represents a single gene. Red points are genes which have significant differential expression (a) or significant differential methylation (b).
